# Alzheimer’s Disease and circadian disruption sex-specifically contribute to a loss of bone maintenance in APP/PS1 model mice

**DOI:** 10.64898/2026.05.01.722089

**Authors:** Noah G. Allen, Carmalena V. Cordi, Joan E. Llabre, Joshua Chuah, Gretchen T. Clark, Angela J. Kubik, Naomi G. Falkenberg, Meaghan S. Jankowski, Rukmani A. Cahill, Ava A. Herzog, Mallika Subash Chander, Deepak Vashishth, Jennifer M. Hurley, Elizabeth A. Blaber

## Abstract

Alzheimer’s Disease and Related Dementias (ADRDs) are linked to reduced bone integrity and increased fracture risk, but the mechanisms that underlie this risk remain poorly defined. Current research suggests that environmental factors, such as diet, sleep, and light exposure can modulate the brain-bone axis, increasing susceptibility to bone loss and fractures. Circadian disruption (CD) associated with ADRDs may exacerbate the effects of disease and aging in the bone. In particular, regulation of bone marrow progenitors may be acutely susceptible to disruption along this axis. Here, we explore the interplay among genetic and environmental factors that influence bone structure, marrow progenitor cell activity, and monocyte-derived macrophages. The APP/PS1 transgenic mouse model (AP) is used as an *in vivo* model of amyloid-beta deposition. High-resolution micro-computed tomography (μCT) identified sex- and genotype-specific responses in trabecular morphometry. Follow-up analysis with Raman spectroscopy (RS) found accumulation of non-enzymatic modifications of the organic matrix and notched three-point bending identified concomitant loss of bone toughness due to both CD and AP. Single-cell RNA sequencing (scRNA-seq) confirmed the presence of oxidative stress signals in the cellular populations of the bone marrow. We further mapped significantly differentially expressed genes (DEGs) from monocytes in the bone marrow to circadian-regulated proteins in monocyte-derived macrophages, revealing dysregulation of circadian timing in macrophages *in vitro*. These findings offer new insights into how environmental disruptions can exacerbate the progression of neurodegenerative disease and bone degradation.

**LAY SUMMARY:** Patients with Alzheimer’s disease have an increased bone fracture risk, but the biological link between brain and bone disease is not well understood. Everyday factors such as altered light exposure (shift work, screens late at night, etc.) can worsen outcomes in the brain and skeleton. Using a mouse model of Alzheimer’s disease, we found that both genetic risk and circadian disruption contribute to weaker bone and altered bone quality. We also identified inflammation and stress responses in bone marrow cells, suggesting that bone marrow may play a key role in linking brain disease to bone fragility.

## INTRODUCTION

Alzheimer’s Disease and Related Dementias (ADRDs) are a group of diseases defined by cognitive decline and memory loss. In concordance with the known cognitive effects of ADRDs, there are also physical comorbidities that worsen patient outcomes. In aging populations, the diagnosis of ADRDs is often associated with the development of osteoporosis.^1–3^ Moreover, bone fracture risk is significantly higher in ADRD individuals with reported hazard ratios ranging from 1.9 to 3.2 compared with age-matched controls, and this fracture risk may be dependent upon the sex of the patient.^4–7^ Despite the wealth of epidemiological data that supports a clinical association between ADRDs and bone health, the underlying mechanisms for this relationship have yet to be elucidated.

Insights into the link between ADRDs and osteoporosis come from understanding the responses of parallel regulatory systems within the brain-bone axis to systemic conditions that contribute to ADRD progression at the tissue level. These systems often share regulatory elements that coordinate physiology across tissues (e.g., Wnt/β-catenin, leptin, estrogen).^8–10^ A key regulatory element across many systems in the brain-bone axis is the circadian clock, which serves as the central pacemaker that synchronizes signaling, metabolism, and cellular function to the day/night cycle.^11^ Inherently, circadian timing is sensitive to environmental cues that organize physiological function across the 24-hour day. For example, light exposure, diet, and exercise all act as exogenous zeitgebers that entrain the central pacemaker (the suprachiasmatic nucleus or SCN), which in turn synchronizes clocks in peripheral tissues, including bone.^12,13^

Mistiming of the circadian clock within the system, termed Circadian Disruption (CD), is triggered by both environmental and disease-related cues and is increasingly recognized as a driver of disease. Notably, CD has been observed in aging populations^13^ and among those afflicted by ADRDs, where reduced sleep quality and duration further amplify circadian dysfunction.^14,15^ In bone tissue, studies have demonstrated that CD can lead to the degradation of trabecular morphology and reduced fracture resistance,^16^ likely driven by clock-regulated control of both osteoblast and osteoclast functions.^17,18^ The inflammatory tone and cellular composition of the bone marrow may be affected by CD due to the sensitivity of mesenchymal and hematopoietic progenitors to environmental stimuli.^19,20^ Moreover, CD has a differential effect based on the sex of the organism, presenting another possible link between the clock, ADRDs, and bone health.^21^ Given the essential role of circadian regulation in both neurological and skeletal homeostasis, we propose that a contributor to the comorbidity between ADRDs and osteoporosis may arise from the disruption of circadian control mechanisms in ADRDs.

Therefore, we examined the effect of CD on the bone marrow microenvironment and the structural integrity of the bone in an ADRD context. In this study, we employed the transgenic APP/PS1 knock-in mouse model (“AP”) in combination with a rotating shift work lighting schedule to induce CD to isolate the effects and interactions of sex, disease, and environment. This model is reported to develop beta-amyloid deposits in the brain as early as 4 to 6 months of age, memory deficits at 6 months,^22^ and behavioral impairments at 12 months.^23^ We assessed the effects of AP and CD at around 8 months as amyloid pathology has begun but neurodegeneration is not yet severe, allowing a view of early-stage disease progression. We assessed the individual and combined effects of the AP genotype and CD exposure on trabecular and cortical bone using µCT, Raman spectroscopy, and fracture mechanics. This analysis was paralleled by single-cell RNA sequencing (scRNA-seq) of the bone marrow and a total proteomics assay of bone marrow-derived macrophages (BMDMs). In total, our model shows that AP and CD drive morphological changes in bone structure that align with conditions of stress and inflammation, extending our understanding of how the brain-bone axis integrates multiple biological regulatory mechanisms that are integral to diseases of aging.

## METHODS

### Animal Model

#### Animal husbandry

Transgenic mice carrying the human amyloid precursor protein with the Swedish mutation and human presenilin (APPswe/PS1, JAX MMRRC Stock #034829) were used as an early-onset Alzheimer’s Disease model. Non-carrier litter mates (LM) were used as control animals. The APPswe/PS1 strain was selected as these animals exhibit amyloid beta deposition in the brain and memory deficits by 6 months of age.^22^ Therefore, we surmised that the collection of tissues around 8 months would allow us to investigate the effects of the early stages of Alzheimer’s Disease.

Animals were maintained in a barrier-controlled facility under an approved IACUC protocol. Mice were group housed (2<n<5) in standard mouse cages with access to water and nutrient bars *ad libitum* for 22 weeks starting at 11 weeks of age. Non circadian disrupted animals were housed under normal 12:12, light:dark conditions. Circadianly disrupted (CD) animals were housed in a rotating light inversion schedule, consisting of four days of a 12:12 light:dark followed by three days of 12:12 dark:light (inverted) pattern, repeated over the course of 22 weeks. Mice were then euthanized at around 8 months of age via cardiac puncture under isoflurane anesthesia and their tissues harvested for this investigation. Exact animal count is reported per assay. Weight and behavior monitoring were used to define humane endpoints and ensure animal health throughout the experiment.

#### Activity monitoring

To determine and validate the effect of the circadianly disruptive conditions on animal behavior, a subset of animals was housed individually in cages equipped with Actimetrics Wireless Low-profile Running Wheels (Model ACT-557-WLP) (females, non-CD/LM: n = 6, AP: n = 7, CD: n = 7, CD/AP: n = 6), or STARR Life Sciences Incage running wheels (male, non-CD/LM: n = 6, AP: n = 6, CD: n = 7, CD/AP: n = 7). Wheel activity was tracked by Philips Respironics VitalView software (version 5.1) and the Actimetrics ClockLab Data Collection System (Model AM1-CD01). Data from week one was not included in the analysis as mice were acclimating to the wheels during that time. Female mice required modified wheel running due to isolation or weight loss. In these cases, affected wheel runners were exchanged with an equivalent experimental animal from group housing. These individually housed (“wheel runner”) animals were not used for tissue analysis if they remained in a wheel running for the full 22-week duration. Using hierarchical linear modeling with nested model comparison, wheel running did not demonstrate significant interaction or additive effects.

### Measurements of Amyloid Beta 42 (Aβ42) from serum and brain tissue homogenates

Brains were dissected, rinsed in 1X PBS, and flash-frozen in liquid nitrogen. One hemisphere was then randomly selected for homogenization. Serum was collected following cardiac puncture and stored at -80^°^C. Full tissue processing details are available in Supplementary Methods. A human Aβ42 ELISA kit (Invitrogen, Cat# KHB3441) was used to quantify Aβ42 levels according to the manufacturer’s instructions. Aβ42 values were normalized to total protein content using the Pierce™ Coomassie Plus (Bradford) Assay Kit (Thermo Fisher, Cat# 23236).

### Micro-Computed Tomography

To investigate morphological changes in cortical and trabecular bone, the right hindlimb was collected and removed of soft tissue before being fixed in a solution of 4% paraformaldehyde for 48 hours, after which it was washed and stored in PBS Ca+/Mg+ until processing. Male LM had an n = 10, AP and CD had an n = 9, and CD/AD had an n = 8. Female LM had an n = 6, AP, CD, CD/AP had an n = 7. Data collection and analysis followed the guidelines established for μCT in rodents.^24^ The hindlimbs were wrapped in PBS Ca- /Mg- soaked gauze and mounted in a Bruker 1272 Skyscan instrument. Male bones were scanned at a 4 µm voxel size, a source voltage of 70 kV, a current of 142 μA, a 0.25 mm aluminum filter, and a 2142 ms exposure time. Due to scheduling limitations, female bones were scanned in a separate batch on the same instrument after hardware maintenance. Thus, female bones were scanned at a 4 µm voxel size, a source voltage 60 kV, a current of 166 μA, a 0.25 mm aluminum filter, and a 1166 ms exposure time. Scans were reconstructed in two batches (males and females) using NRecon (version 1.7.4.6, Bruker) with Gaussian smoothing kernel = 2, ring artifact correction = 10, and beam hardening correction = 15% (males) and 49% (females). Post-scan alignment correction was conducted for each sample separately.

Reconstructed scans were rotated for the volume of interest (VOI) assignment in the femoral head, mid-shaft femur, and distal femur with transverse cutting planes perpendicular to the direction of weight loading specific to each region (DataViewer version 1.5.6.2, Bruker). Femoral head VOI: trabecular region 5 slices (0.02 mm) distal of the growth plate extending for 25 slices (0.1 mm, females) or 50 slices (0.2 mm, males) since the male bones were slightly larger. Mid-shaft cortical VOI: a 0.24 mm thick section centered at the femoral mid-point, as measured from the most proximal section of the femoral head to the end of the distal epiphysis. Distal femur VOI: a 1.12 mm thick section 0.02 mm proximal to the growth plate. Trabecular and cortical volumes were defined by automated segmentation applied in CTAn (version 1.18.8.0, Bruker) using procedural binary operations (open, dilate, close, and erode). Parameters were calculated using the 2D and 3D analysis algorithms provided in CTAn. Complete instrument, reconstruction, and segmentation details can be found in the Supplementary Methods.

### Biomechanical Characterization of Bones

The biomechanical properties of bone were characterized using fracture mechanics-based tests for toughness, which provide a measure of bone’s resistance to fracture due to a pre-existing flaw.^25,26^ In a separate cohort of mice that underwent the same lighting protocols, we randomly selected one left or right femur per mouse from each lighting. Male LM, AP, and CD had an n=4 and CD/AP had an n = 5. Female LM had an n = 4, and AP, CD, and CD/AP had an n=5. The protocol for the sample preparation of femora has been described in detail previously.^16,27^ Briefly, bones were notched at the mid-diaphyseal region and scanned with µCT prior to three-point bending using an electromechanical testing system (EnduraTEC 3200, TA Instruments, New Castle, DE).Full details are available in the Supplemental Methods.

### Raman spectroscopy (RS) for Assessing Bone Matrix Composition

Raman spectroscopy (RS) measures the wavelength shift of scattered light to describe the composition of bone tissue in terms of its mineral and organic compartments.^28^ Here, we followed established protocols to analyze the bone composition using RS.^27,29^ Male LM and AP had an n = 4, CD had an n = 3, and CD/AP had an n = 5. Females (LM, AP, CD, CD/AP) had an n = 5. Full details available in Supplementary Methods.

### Quantification of Fluorescent Advanced Glycation End-Products (fAGEs) in Bone Matrix

Briefly, protein isolates from midshaft femur samples were extracted for both a quinine sulphate fluorescence assay and a collagen content assay. The fAGEs content for each sample was then calculated and expressed as ng of quinine per mg of collagen as described previously.^16,30^ Full details available in Supplementary Methods.

### Bone Marrow Single Cell Analysis

Bone marrow was collected from the left femur of male mice (n = 1 per group) by removing soft tissue and cutting the proximal and distal epiphyses. Femurs were flushed with cryo-protectant media (44.5% DMEM- α, 44.5% FBS, 10% DMSO, 1% Penicillin/Streptomycin) into cryovials which were sealed on dry ice and stored at -80^°^C. Following 10X Genomics recommendations, 8,000 live cells were targeted for recovery. Library preparation and sequencing were performed by NovoGene (Sacramento, CA) at a depth of approximately 20,000 reads per cell. Raw FASTQ were processed and aligned using the 10X Genomics Cloud Analysis Platform running CellRanger (v7.1.0) and aligned to the *Mus musculus* reference genome GRCm38 (GCF_000001635.20). Full sample handling details are available in Supplementary Methods.

Count, barcode, and feature matrices were merged in R into a single dataset with treatment metadata retained to allow normalization across all cells. Data processing was performed using Seurat (v5.2.1).^31^ Cells were filtered by feature counts (100-5,500 genes) and mitochondrial transcript content (< 7.5%). Expression was normalized and scaled using the Seurat functions, NormalizeData and ScaleData. Gene ontology analysis was conducted using the R packages clusterProfiler (v4.14.4) and gprofiler2 (v0.2.3).

### Bone Marrow Derived Macrophage Proteomics for Circadian Assays

#### BMDM Isolation, Synchronization, and Analysis

A subset of animals was used for the isolation of bone marrow to generate BMDMs. Femora and tibiae were collected immediately following euthanasia. As defined in established protocols,^32^ bone marrow was flushed, and macrophages were differentiated in MCSF-supplemented media. Cells were synchronized using a serum shock protocol. At 16 hours post-shock (PS), cells were washed, lifted, and preserved as a frozen pellet. Sampling continued every three hours until PS time 40. This was repeated in three cohorts of mice (n = 2, per cohort) from each experimental condition.

Frozen cell pellets were processed at the Thermo Fisher Center for Multiplexed Proteomics at Harvard Medical School. Briefly, proteins were extracted, reduced, alkylated, exposed to trypsin digestion, and labeled using TMTpro-18plex reagents. Peptides were desalted, fractionated, and analyzed by LC-MS/MS. Peptide identification was performed from MS2 spectra and mapped to the *Mus musculus* reference proteome (UniProt ID UP000000589).

Rhythmicity in protein levels was assessed using the Extended Circadian Harmonic Oscillatory (ECHO) model (v4),^33^ which fits expression time series data to detect circadian patterns. Circadianly expressed proteins were defined by the ECHO model statistics that met the following criteria: (a) rhythmic profile consistent with harmonic, forced, or damped oscillations, (b) period length between 20–28 hours, and (c) Benjamini–Hochberg adjusted p < 0.05. More details on the data acquisition can be found in the Supplementary Methods and in the associated data set from Cordi, et al. (2026).^34^

### Statistical Approach

Statistical tests and plotting were conducted in Graph Pad Prism (v10). Each data set was subjected to normality testing (Shapiro-Wilkes) and outlier testing (Grubb’s Iterative). Outliers were removed from statistical analysis and reported on plots as red points. Two-way ANOVA was applied to test the AP genotype effect and CD effects within each sex separately. Post-hoc Tukey’s pairwise comparisons are reported in figures (* p<0.05, **p<0.01, ***p<0.001, ****p<0.0001). No statistical method was used to predetermine the sample size; sample size was determined based on literature and previous experience to achieve biologically important differences. The investigators were not blinded during experiments. Animals were randomly assigned to non-CD and CD lighting groups.

#### Activity monitoring metrics

Aging was considered using a repeated measures mixed-effects analysis to compare age (weeks 2 and 3 to weeks 21 and 22) and experimental condition.

#### Aβ42 ELISA

Aβ42 ELISA on brain homogenate in non-AP carriers was zero-inflated and failed statistical assumptions, therefore we used the assay limit of detection (LOD, 15.6 pg/mL) to perform contingency analysis by Fisher’s Exact test.

#### Bone marrow single cell analysis metrics

Single cell RNA sequencing was conducted on one male non-wheel runner animal randomly selected from each experimental condition. This single replicate, cell-level, differential expression approach provides an exploratory representation, though is limited in the ability to model biological variability. Single cell analysis was conducted in Seurat (v5.2.1). Heatmap plotting utilized pheatmap (v1.0.12). Single cell pairwise differential testing used the Model-based Analysis of Single-cell Transcriptomics package (MAST, v1.27.1)^35^ to compute Log2FC and Bonferroni-Hochberg adjusted p-values. The Scanpro package (v0.3.2)^36^ was used to estimate cell proportions.

## RESULTS

### AD-related activity decline is ameliorated by CD in females

Given the strong links between CD, ADRDs, and bone health, we hypothesized that exposing mice with an ADRD background to a CD protocol would enhance the negative bone health effects noted in patients with ADRDs. To validate our model, amyloid beta load was assessed in serum and brain tissue collected at the experimental endpoint. A significant increase in total Aβ42 was detected in both the serum (**Supplemental Figure 1 A, B**) and the brain tissue homogenate (**Supplemental Figure 1 C, D**) of AP genotype animals (**Supplemental Table 1**). A detailed interpretation of brain region-specific plaque formation is available elsewhere.^34^ To assess the effects of CD, running wheels were added to a subset of cages. Interdaily Stability (IS), a metric of rhythm consistency between days, was reduced in animals exposed to CD, regardless of sex or genotype, by the end of the 22-week protocol (**Figure 1 A, B**), confirming successful disruption of circadian rhythms. Intradaily Variability (IV), which reflects fragmentation of activity and rest periods, was not increased (**Figure 1 C, D**). This suggests that while day-to-day rhythmicity was disrupted, patterns within a day remained intact.

**Table 1.** Distal femora trabecular morphology assessed by µCT imaging. Results shown are mean and standard deviation for each group and parameter. Two-way ANOVA results are shown. Fstat = calculated F statistic, p.val = p value calculated. DS = Dayshift, CD = circadian disruption, LM = Littermate, AP = APP/PS1 transgenic, Conn.D = Connectivity density, DA = Degree of anisotropy, SMI = structural model index, Tb.N = Trabecular number, Tb.Pf = Trabecular pattern factor, Tb.Sp = Trabecular separation, Tb.Th = Trabecular thickness. Bolded red values highlight significant 2-way ANOVA result (p<0.05).

| Distal Femur |  | Condition | BV/TV% | Conn.D<br>(1/mm<sup>3</sup>) | DA | SMI | Tb.N<br>(1/mm) | Tb.Pf<br>(1/mm) | Tb.Sp<br>(mm) | Tb.Th<br>(mm) |
| --- | --- | --- | --- | --- | --- | --- | --- | --- | --- | --- |
| Mean±SD | Male | DS/LM | 11.93<br>$\pm 4.918$ | 260.975<br>$\pm 116.025$ | 1.657<br>$\pm 0.216$ | 1.386<br>$\pm 0.33$ | 2.209<br>$\pm 0.905$ | 15.744<br>$\pm 3.981$ | 0.302<br>$\pm 0.072$ | 0.055<br>$\pm 0.002$ |
| | | DS/AP | 9.506<br>$\pm 2.659$ | 194.605<br>$\pm 57.66$ | 1.655<br>$\pm 0.212$ | 1.916<br>$\pm 0.264$ | 1.929<br>$\pm 0.542$ | 24.791<br>$\pm 4.468$ | 0.269<br>$\pm 0.043$ | 0.049<br>$\pm 0.004$ |
| | | CD/LM | 12.591<br>$\pm 2.765$ | 371.936<br>$\pm 154.45$ | 1.501<br>$\pm 0.246$ | 1.156<br>$\pm 0.625$ | 2.731<br>$\pm 0.661$ | 14.863<br>$\pm 8.187$ | 0.239<br>$\pm 0.017$ | 0.047<br>$\pm 0.005$ |
| | | CD/AP | 11.735<br>$\pm 4.999$ | 243.756<br>$\pm 103.151$ | 1.549<br>$\pm 0.197$ | 1.88<br>$\pm 0.593$ | 2.457<br>$\pm 1.244$ | 20.358<br>$\pm 13.472$ | 0.221<br>$\pm 0.009$ | 0.05<br>$\pm 0.007$ |
| 2-way ANOVA | Lighting | Fstat | 1.170 | 4.246 | 3.194 | 0.670 | 3.285 | 0.912 | 10.294 | 4.434 |
|  |  | p.val | 0.287 | **0.048** | 0.083 | 0.419 | 0.079 | 0.347 | **0.003** | **0.044** |
|  | Genotype | Fstat | 1.509 | 6.269 | 0.098 | 14.840 | 0.917 | 6.834 | 2.190 | 0.430 |
|  |  | p.val | 0.228 | **0.018** | 0.757 | **0.001** | 0.346 | **0.014** | 0.150 | 0.517 |
|  | Interaction | Fstat | 0.345 | 0.633 | 0.118 | 0.355 | 0.000 | 0.408 | 0.174 | 6.771 |
|  |  | p.val | 0.561 | 0.432 | 0.733 | 0.556 | 0.992 | 0.528 | 0.679 | **0.014** |
| Mean±SD | Female | DS/LM | 5.056<br>$\pm 4.002$ | 95.426<br>$\pm 42.105$ | 1.44<br>$\pm 0.165$ | 2.919<br>$\pm 0.607$ | 0.884<br>$\pm 0.65$ | 41.978<br>$\pm 14.415$ | 0.446<br>$\pm 0.104$ | 0.054<br>$\pm 0.007$ |
| | | DS/AP | 6.834<br>$\pm 2.768$ | 265.772<br>$\pm 94.583$ | 1.454<br>$\pm 0.22$ | 2.503<br>$\pm 0.268$ | 1.162<br>$\pm 0.445$ | 30.728<br>$\pm 5.526$ | 0.405<br>$\pm 0.062$ | 0.059<br>$\pm 0.004$ |
| | | CD/LM | 4.239<br>$\pm 3.138$ | 111<br>$\pm 57.221$ | 1.523<br>$\pm 0.18$ | 3.154<br>$\pm 0.463$ | 0.683<br>$\pm 0.499$ | 42.308<br>$\pm 14.396$ | 0.502<br>$\pm 0.12$ | 0.06<br>$\pm 0.011$ |
| | | CD/AP | 7.218<br>$\pm 4.518$ | 337.599<br>$\pm 210.718$ | 1.388<br>$\pm 0.194$ | 2.296<br>$\pm 0.151$ | 1.242<br>$\pm 0.701$ | 30.031<br>$\pm 5.664$ | 0.4 $\pm 0.09$ | 0.057<br>$\pm 0.005$ |
| 2-way ANOVA | Lighting | Fstat | 0.023 | 0.685 | 0.013 | 0.007 | 0.074 | 0.002 | 0.477 | 0.491 |
|  |  | p.val | 0.880 | 0.417 | 0.909 | 0.932 | 0.788 | 0.966 | 0.497 | 0.490 |
|  | Genotype | Fstat | 2.843 | 14.132 | 0.605 | 14.726 | 3.501 | 7.487 | 3.720 | 0.034 |
|  |  | p.val | 0.105 | **0.001** | 0.445 | **0.001** | 0.074 | **0.012** | 0.066 | 0.856 |
|  | Interaction | Fstat | 0.181 | 0.284 | 0.920 | 1.777 | 0.394 | 0.014 | 0.673 | 1.695 |
|  |  | p.val | 0.674 | 0.600 | 0.348 | 0.197 | 0.536 | 0.906 | 0.420 | 0.206 |

**Figure 1.**
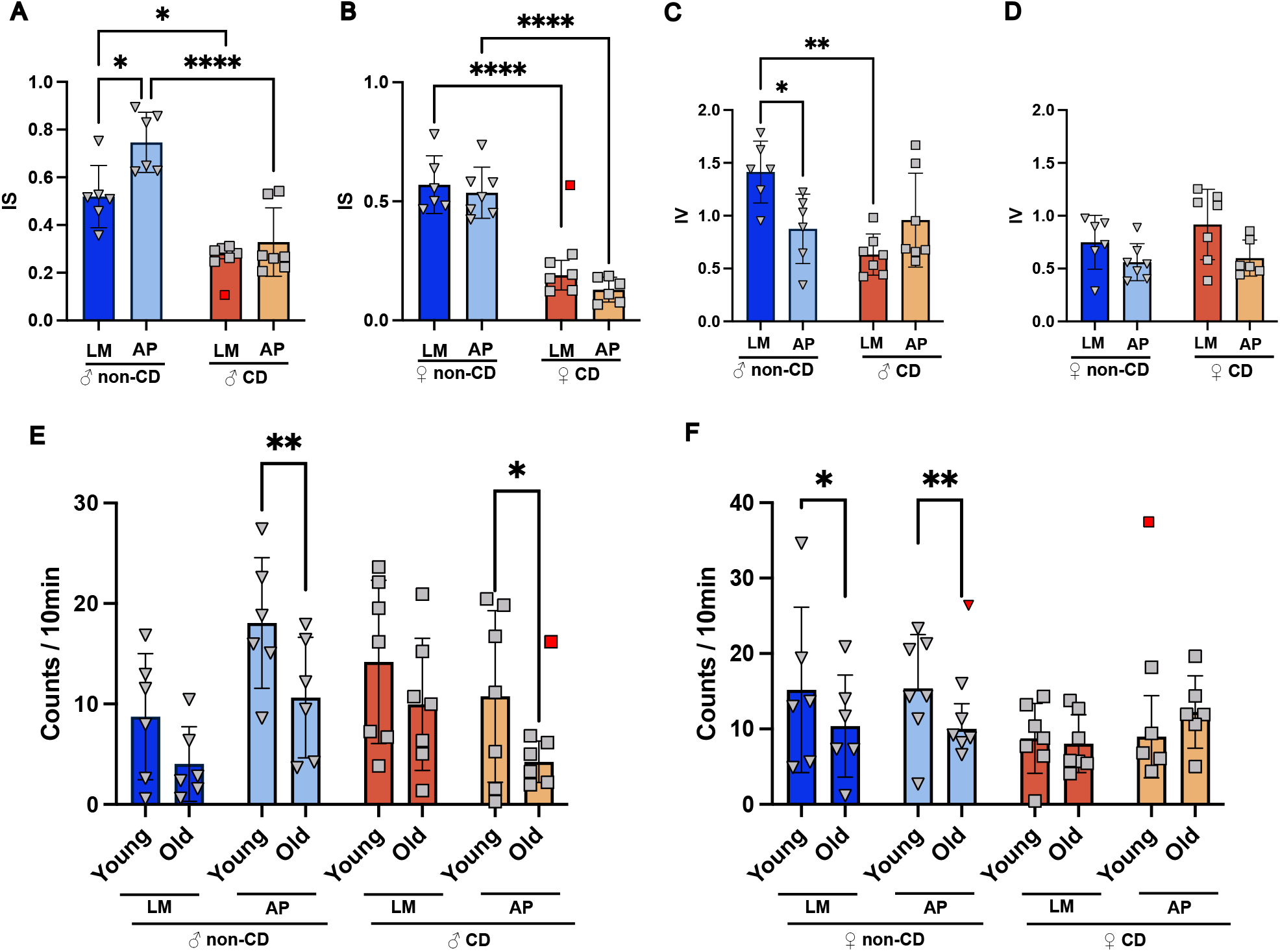
Age, sex, genotype, and lighting differentially affect murine activity. **A)** Male Interdaily Stability (IS) and **B)** Female IS behavioral patterns during the last two weeks of the experiment. **C**) Male Intradaily Variability (IV) and **D)** Female IV behavioral patterns during the last two weeks of the experiment. **E)** Activity volume (counts per 10-minute interval) in male and **F)** female mice. Data from the first two weeks of the experiment are referred to as young (10 weeks) whereas data from the last two weeks are referred to as old (32 weeks). Tukey’s post-hoc pairwise comparison displayed (* p<0.05, **p<0.01, ***p<0.001, ****p<0.0001). Dark blue = control/littermates (LM), Light blue = Alzheimer’s genotype (AP), Dark orange = Circadianly disrupted (CD), and Light orange = CD/AP.

Given that our CD protocol successfully modulated organismal rhythms, we next compared the rotation counts at the beginning and end of the experiment to assess changes in activity over time due to CD. We found that, in general, activity decreased with age in male LM and AP mice, regardless of CD. A repeated-measures analysis demonstrated male activity was significantly affected by age (**Supplemental Table 2**). This trend was stronger in the AP genotype, as Tukey’s pairwise testing found a significant decrease in activity in male mice with the AP genotype, suggesting the AP genotype exacerbated the effect of age in this model (**Figure 1 E**). This trend was similar in females, however repeated-measures analysis detected a significant interaction effect (**Supplemental Table 2**). Tukey’s pairwise testing found a significant age-related decrease in activity in female mice without CD (**Figure 1 F**). The female AP mice that underwent CD did not show a decrease in activity, but rather a non-significant increase in activity (**Figure 1 F**), demonstrating a combined effect of CD and AP in females. Of note, there was no associated change in body mass, liver weight to body mass ratio, or liver lipids (**Supplemental Figure 2**), suggesting that our CD model did not induce fundamental metabolic changes.

### AP genotype drives negative trabecular changes in male mice

Given the known individual effects of CD and ADRDs on bone health, we next analyzed the femora from mice that underwent our tailored lighting protocol to determine if CD and AP interact in the bone. We used μCT reconstructed scans to evaluate the femoral head, cortical mid-shaft, and distal femur, as the femur is often remodeled during disease and stress.^8,20^ Given the noted technical differences (x-ray voltage and exposure time) and observed sexual dimorphism, we analyzed the effect of AP and CD on males and females independently to understand their effect on each sex.

A two-way ANOVA was performed on genotype (LM, AP) and lighting (non-CD, CD) conditions within each sex. In male mice, a significant genotype effect was observed, as AP was associated with decreased connectivity density (Conn.D), increased structural model index (SMI), and increased trabecular pattern factor (Tb.Pf) (**Table 1**). Tukey’s pairwise comparison supported the significant effect of AP on Tb.Pf (**Supplemental Figure 3 C**). Among CD-treated males, SMI was significantly increased in AP compared to LM animals (**Figure 2 B**). The AP-related increase in Tb.Pf and decrease in Conn.D indicate a sparse network,^24,37^ while increased SMI relates to a thinner more rod-like trabecular structure.^24^ These findings prompted further analysis of trabecular microarchitecture in the femoral head to assess whether genotype effects were consistent across anatomical regions. Trabeculae in the femoral head showed a significant genotype effect where AP increased both SMI and Tb.Pf (**Supplemental Table 4** and **Supplemental Figure 4**) and decreased degree of anisotropy (DA), again indicating a thinner, more disorganized trabecular network. Taken together, trabeculae in both the femoral head and distal femur suggest that the AP genotype results in a less dense network of trabeculae, indicating altered structural integrity not attributable to differences in activity (**Figure 1 F**).

### Trabecular thickness in the male distal femur shows an AP-dependent response to CD

As noted previously, the interactions of CD and AP may be important for the outcomes of bone health. A two-way ANOVA in males found a significant interaction between CD and AP in trabecular thickness (Tb.Th) and the Tukey’s test revealed that trabeculae in the CD group were significantly thinner compared to LM controls (**Figure 2 F**), suggesting that the effect of CD is modulated in the presence of AP in males. While our two-way ANOVA of the distal femur demonstrated that the primary driver of change was the AP genotype, this data demonstrates there is a potential effect of CD on the trabeculae, indicating an overall negative effect under the combination of CD and AP on the bone health of males.

### AP does not affect distal trabeculae but interacts with CD in the femoral head of female mice

In the distal femur of female mice, while genotype had an effect, the directionality of the change was different compared to males. Two-way ANOVA found a significant AP genotype effect, with increased Conn.D and reduced SMI (**Table 1, Figure 2**). This was supported by the Tukey’s analysis, which showed a significant AP effect, though only in the CD treated group (**Figure 2 D and G**). These metrics demonstrate that CD/AP females had a denser and more interconnected trabecular network. Although this phenotype contrasts with the trabecular deterioration observed in males, it has also been observed in the 5xFAD model,^38^ and likely reflects sex- and age-dependent differences in bone remodeling. Considering the pre-menopause state at around 8-months of age, elevated estrogen levels may preserve trabecular microarchitecture, modifying the skeletal impact of AP and CD at this time point. We again looked to the femoral head to compare different anatomical trabecular regions and found that AP females showed a significant decrease in Tb.Sp (**Supplemental Table 4** and **Supplemental Figure 5 G**). Additionally, two-way ANOVA identified significant lighting and genotype interactions in trabecular microarchitecture parameters (DA and SMI, **Supplemental Table 4** and **Supplemental Figure 5 C, D**) indicating altered structural organization rather than net bone gain or loss.

**Figure 2.**
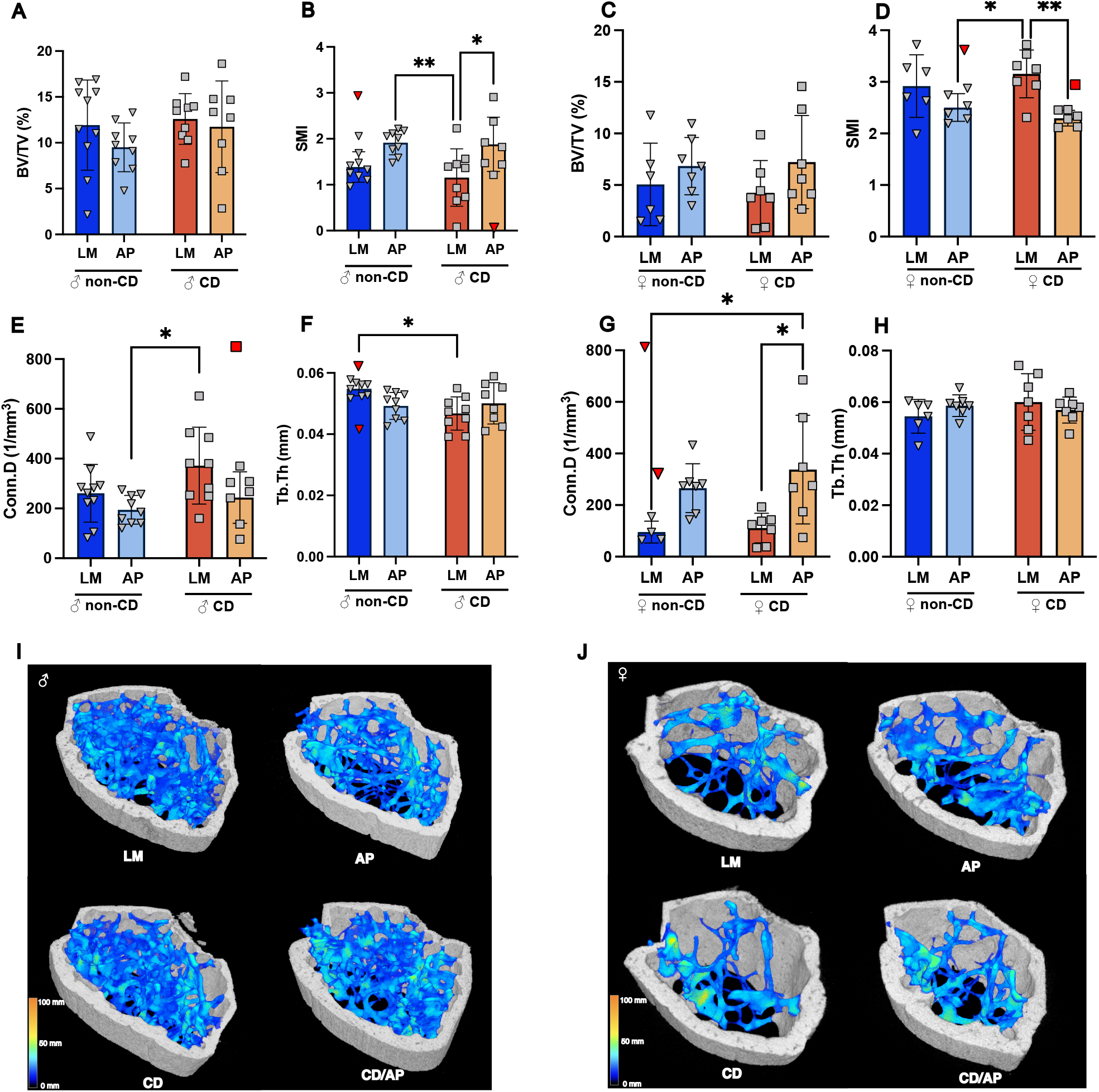
The distal femur is sex-specifically affected by the AD genotype. **A)** Male bone volume fraction (BV/TV%) by genotype and lighting condition. **B)** Male structural model index (SMI) by genotype and lighting condition. **C)** Female bone volume fraction (BV/TV%) by genotype and lighting condition. **D)** Female structural model index (SMI) by genotype and lighting condition. **E)** Male connective density (Conn.D) by genotype and lighting condition. **F)** Male trabecular thickness (Tb.Th) by genotype and lighting condition. **G)** Female connective density (Conn.D) by genotype and lighting condition. **H)** Female trabecular thickness (Tb.Th) by genotype and lighting condition. Shiftwork conditions are labeled in orange and standard lighting conditions are labeled in blue **I)** Representative images of Male and **J**) Female distal femur reconstructions from micro-computed tomography (µCT). Analysis is conducted on the trabecular bone (blue) contained within the cortical bone (white). Dark blue indicates thinner trabeculae and yellow indicates thicker trabeculae. Tukey’s post-hoc pairwise comparison displayed (* p<0.05, **p<0.01, ***p<0.001, ****p<0.0001). Red points indicate outlier removed from analysis as determined by Grubb’s test. Dark blue = control/littermates (LM), Light blue = Alzheimer’s genotype (AP), Dark orange = Circadianly disrupted (CD), and Light orange = CD/AP.

### CD and AP increased Carboxymethyl-lysine (CML) in the bone matrix, affecting toughness without impacting mineral phase or cortical morphometry

Given that trabecular bone is highly dynamic, and skeletal deficits associated with CD and AP may accumulate over time, we next assessed cortical bone as a long-term indicator of chronic effects.^39^ Femoral midshaft µCT analysis of cortical bone in males and females revealed no changes in total cross-sectional area (Tt.Ar), cortical bone area (Ct.Ar), cortical thickness (Ct.Th), endocortical (Ec.Pm), nor periosteal perimeter (Ps.Pm) between genotypes or lighting conditions (**Supplemental Table 5** and **Supplemental Figure 6**). The absence of detectable morphological changes in cortical bone prompted us to explore cortical bone composition using Raman spectroscopy (RS), as changes in bone quality can precede detectable losses in bone mass.^28^ RS analysis indicated that compositional differences were restricted to the organic phase of the bone. Specifically, the AP genotype exhibited increased accumulation of carboxymethyl-lysine (CML), a marker of collagen glycation, in both males (**Figure 3 A**) and females (**Figure 3 B**). However, mineral composition (crystallinity, carbonate to phosphate ratio, and mineral to matrix ratio) showed no effect of genotype or lighting condition (**Supplemental Figure 7** and **Supplemental Table 5**). Quantification of fluorescent advanced glycation end-products (fAGEs) showed a parallel trend with CML accumulation, with the most pronounced changes in females (**Supplemental Figure 7 J** and **Supplemental Table 5**). In total, the compositional makeup of the bones suggests that AP, not CD, is the primary driver of oxidative stress-based modifications.

**Figure 3.**
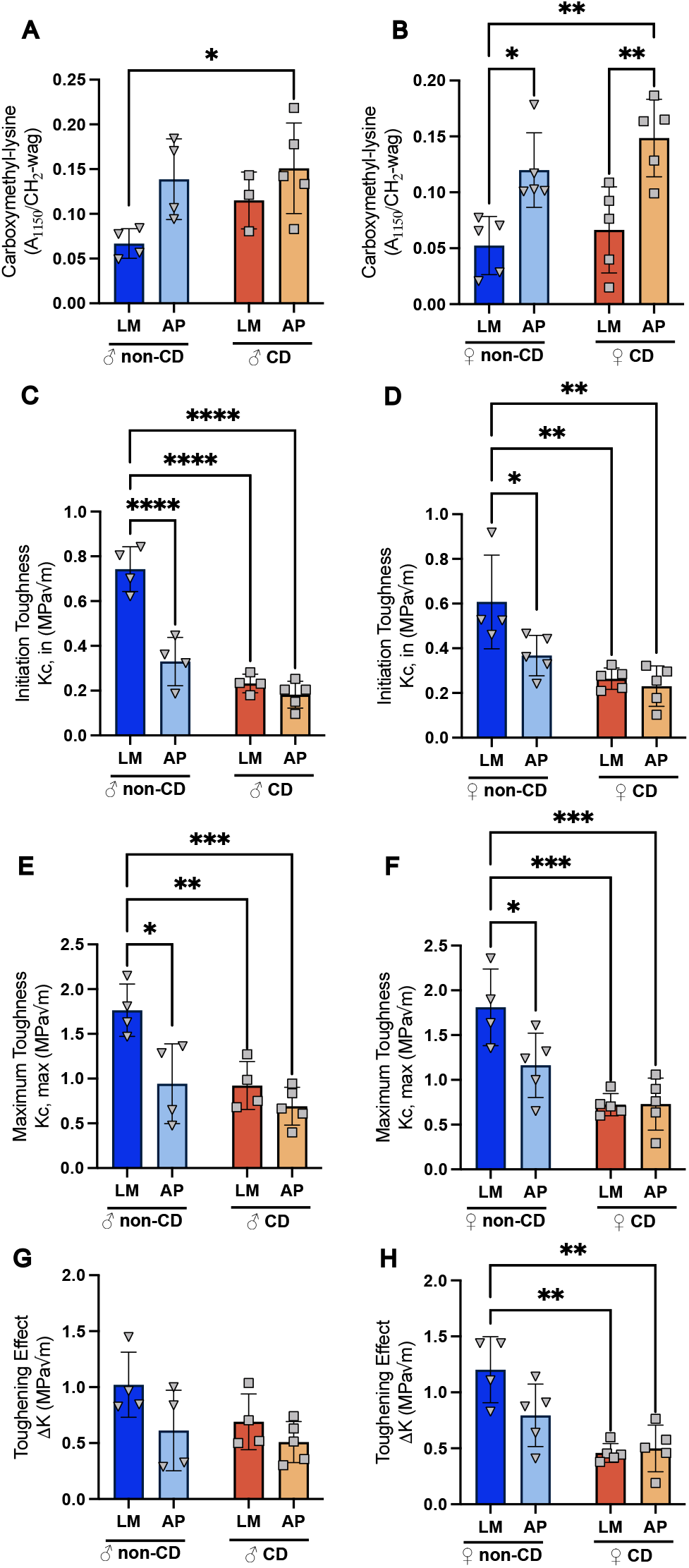
Organic phase Raman spectroscopy and mechanical testing. Male **A**) and Female **B**) Carboxylmethyl-lysine peak in Raman spectroscopy. Male **C**) and Female **D**) Initiation toughness from notched three-point bending strength testing. Male **E)** and Female **F)** maximum toughness. Male **G)** and **Female H)** toughening effect derived from initiation and maximum toughness. Tukey’s post-hoc pairwise comparison displayed (* p<0.05, **p<0.01, ***p<0.001, ****p<0.0001). Dark blue = control/littermates (LM), Light blue = Alzheimer’s genotype (AP), Dark orange = Circadianly disrupted (CD), and Light orange = CD/AP.

Because degraded organic phase quality via crosslinking modification compromises the bone’s ability to absorb energy, we next performed notched three point bending biomechanical toughness tests on the femora.

In both males and females, initiation toughness (the energy required to induce fracture) was significantly reduced by both CD and AP, as detected by two-way ANOVA and post-hoc pairwise Tukey’s analysis (**Supplemental Table 6, Figure 3 C, D**). Similarly, maximum toughness (the total energy absorbed by the bone) was significantly reduced by both CD and AP in both sexes, supported by post-hoc analysis (**Supplemental Table 6, Figure 3 E, F**). However, toughening effect (difference in initiation and maximum toughness) uncovered sex specific effects. Male mice had a mild genotype effect (**Supplemental Table 6**), though not significant in the post-hoc testing. Female mice demonstrated a significant lighting and interaction effect (**Supplemental Table 6**), post-hoc testing indicated a significant decrease due to CD (**Figure 3 H**). This suggests that while both AP and CD reduced the intrinsic resistance to crack initiation in males and females, CD specifically impaired extrinsic toughening in females, indicating reduced capacity for energy dissipation.^26^

### Single cell sequencing of CD-exposed bone marrow indicates a response to stress in the AP condition for male mice

Given the observed changes in bone quality, particularly in male mice, we next sought to determine underlying mechanisms. Because bone marrow progenitor cells and their transcriptional programs are sensitive to stimuli that influence maintenance of bone mineral density and bone volume,^20,40^ we performed single-cell RNA-sequencing (scRNA-seq) of bone marrow from male mice (**Figure 4A**). Bulk differential expression analysis using a likelihood ratio test (LRT) identified several condition specific genes. Clustering revealed that cluster 6 (**Figure 4 B**) showed increased expression in both the AP and CD conditions and had significant GO enrichment in terms related to innate immune, stress, external stimulus, and antioxidant response pathways (**Supplemental Table 7**).

To investigate antioxidant mechanisms, we highlighted common antioxidant transcripts and clustered them by similarity in expression (**Figure 4 C**). Pairwise comparison testing using the MAST test found that glutathione-disulfide reductase (*Gsr*) was significantly up-regulated in both the non-disrupted AP group and the CD/AP group compared to the control (**Supplemental Table 8**). As *Gsr* contributes to the pool of reducible glutathione, upregulation of its transcript is a direct consequence of increased oxidative stress and thus is a marker of inflammatory stress.^41^ In comparison, expression of antioxidant genes in the CD condition are unperturbed and closely resemble the control condition (**Figure 4 C, Supplemental Table 8)**. These findings suggest that while AP contributes to oxidative stress, CD alone may be insufficient to trigger a broad activation of antioxidant response in the bone marrow. This is consistent with our earlier observation that AP had a pronounced negative effect on bone morphometry.

### CD and AP differentially affect monocyte and osteoblast population proportions in male mice

The activation, recruitment, and differentiation of different bone marrow cell sub-types represent a critical axis of bone homeostasis, which may be disrupted in AP and CD conditions. A UMAP projection of the scRNA-seq data from male mice predicted 15 primary cell type clusters (**Figure 4 D** and **E**), though the number of cells in each group shifted depending on condition, suggesting a combined effect of CD and AP on bone marrow cell type distributions (**Figure 4 D**). Using bootstrapped estimates, we found the proportion of monocytes in the bone marrow decreased significantly in the AP genotype, though CD alleviated this effect (**Figure 4 F**). In contrast, CD suppressed the osteoblast proportion (**Figure 4 G**) with significant reductions observed in both LM mice (control LM vs CD) and AP mice (control AP vs CD/AP). Given the sensitivity of monocytes to peripheral signals and their role in the osteoclast lineage,^20^ we decided to focus our analysis on this population.

**Figure 4.**
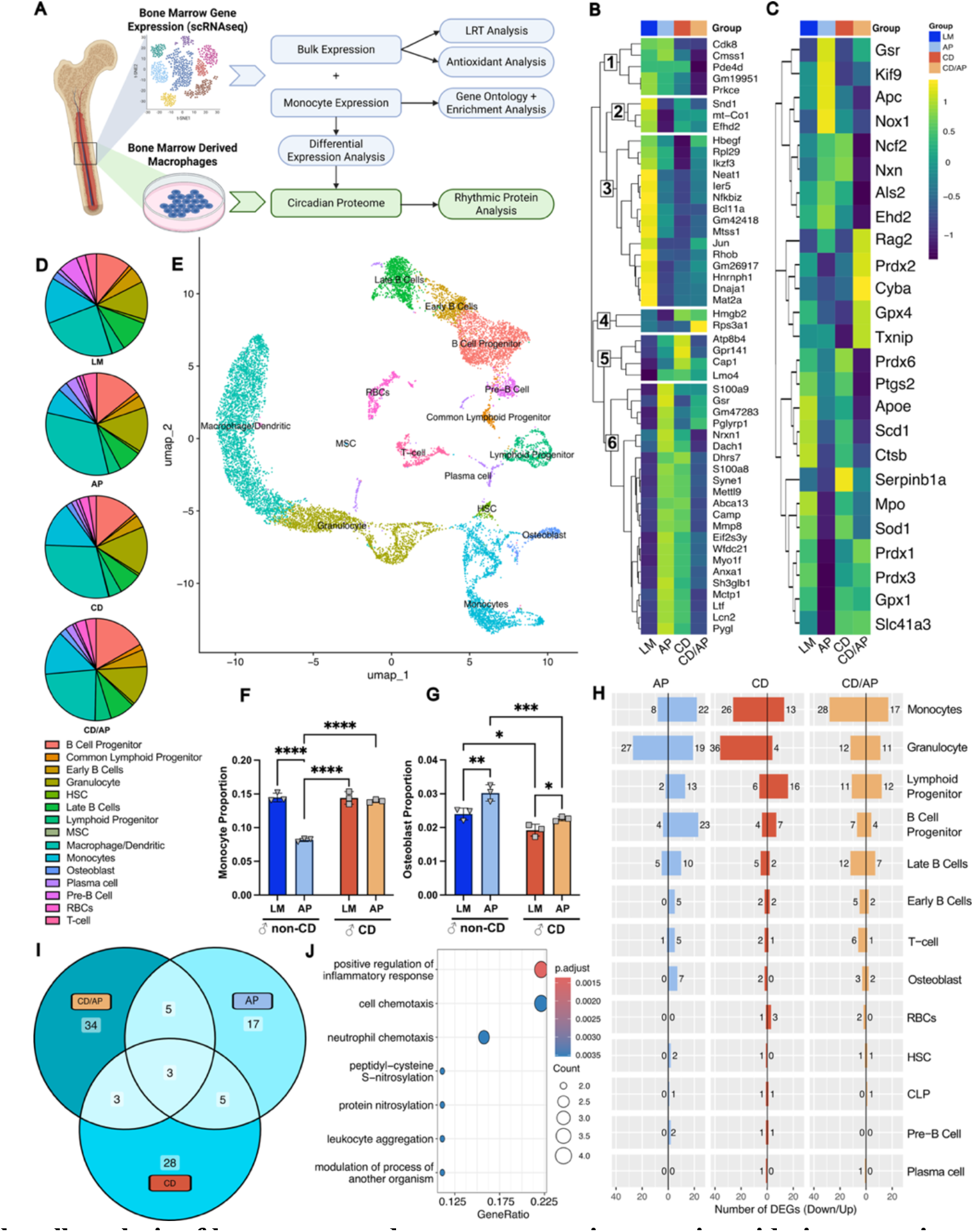
Single cell analysis of bone marrow demonstrates an increase in oxidative stress in response to AP. **A)** Bone marrow single-cell RNA analysis steps; differentially expressed genes from monocytes were used to filter the circadian proteome (Figure 5). **B)** The 51 top differentially expressed genes (measured by LRT, adjusted p-value <1×10^-7^). Clusters were computed and represented in a heatmap with hierarchical clustering. **C)** A heat map of the differential expression of antioxidant genes known to moderate oxidative stress. **D)** Cell subtype proportions according to single cell Seurat analysis. **E)** UMAP projection of individual cells and clustering based on shared cell type expression from Seurat analysis, representative of all cells from each condition (Male: LM, AP, CD, CD/AP). **F)** Monocyte and **G)** osteoblast proportions calculated via bootstrapping to simulate sampling variability. Tukey’s post-hoc pairwise comparison displayed (* p<0.05, **p<0.01, ***p<0.001, ****p<0.0001). **H)** Differential expression using the Seurat function FindAllMarkers with the MAST test. **I)** Number of specific and shared DEGs per condition in monocytic cells. **J)** Gene Ontology Biological Process terms unique to DEGs in CD condition. Dark blue = control/littermates (LM), Light blue = Alzheimer’s genotype (AP), Dark orange = Circadianly disrupted (CD), and Light orange = CD/AP.

### Monocyte inflammation and oxidative stress mark divergent responses to CD and AP in male mice

We next performed differential expression analysis of male scRNA-seq data using the MAST test to identify cell-type-specific responses to environment and genotype. We found that monocytes and granulocytes had the largest number of differentially expressed genes (DEGs) (**Figure 4 H**), and DEG sets appeared highly cell type–specific (**Supplemental Figure 8 C-E**). Monocyte DEGs were largely condition-specific with few shared between conditions (**Figure 4 I** and **Supplemental Table 8**). DEGs in the CD condition found significant GO term enrichment for “response to lipopolysaccharide” and “leukocyte migration” (**Figure 4 J, Supplemental Table 9**). This was driven by up-regulation of the *S100a8* and *S100a9*, common biomarkers in rheumatoid arthritis,^42^ suggesting CD stimulates inflammatory immune pathways in bone marrow-resident monocytes. In contrast, AP monocytes demonstrated GO term enrichment for cellular maintenance processes (**Supplemental Table 9**). Notably, AP monocytes uniquely up-regulated *Gsr* (**Supplemental Figure 8 B)**, a signal of oxidative stress.^41^ Meanwhile, CD/AP DEGs showed enrichment of GO terms related to negative regulation of phosphorylation and MAPK cascade (**Supplemental Table 9**), supported by significant down-regulation of the Dual Specificity Phosphatase 1 (*Dusp1)* transcript, a regulator in inflammation and the MAPK pathway.^43^ Together this suggests that the co-occurrence of CD and AP may heighten both inflammatory and stress responses through loss of regulatory control. To determine whether these transcriptional changes reflect disease-relevant AD processes, we performed Gene Set Enrichment Analysis (GSEA) using a reference dataset^44^ containing both human patient data and mouse amyloid models. Monocytes in the AP and AP/CD conditions were enriched for AD-associated microglial signatures, including, disease-associated microglia (DAM) and tau associated microglia, whereas CD alone showed no enrichment (**Supplemental Figure 8 A**). To determine if these transcriptional alterations persist in differentiated immune lineages, we next examined bone marrow-derived macrophages.

### Bone marrow-derived macrophages *in vitro* demonstrate loss of pathway coordination in circadian timing

Previously, we have demonstrated that extensive circadian coordination of the proteome of bone marrow-derived macrophages from male mice can regulate oxidative stress responses.^32,45^ Here, we used a similar assay to examine differences in oscillating proteins in BMDMs from CD and AP male and female mice (**Figure 5 A**). Filtered circadian proteins are exemplified in **Figure 5 B**, while a full analysis is reported elsewhere.^34^ Here we specifically focused on proteins also identified as DEGs in monocytes from our male scRNA-seq dataset (**Supplemental Figure 8 B**). Of the 95 DEGs identified in monocytes, 12 were found in the circadian proteome of the BMDMs. After aligning proteomic time-series data to circadian time (CT) using ECHO, rhythmic proteins in controls were classified as early-or late-peaking relative to CT12, and their peak timing was subsequently evaluated in the experimental condition. In males, proteins that peaked early in the circadian day under LM conditions showed a coordinated phase delay in the CD and AP condition (**Figure 5 C**). In females, this coordination was not maintained, but there were significant changes in the phase of the DEG proteins (**Figure 5 D**). In total, these data suggest CD and AP disrupt key proteins shared between monocyte and macrophage populations, reflecting a stress response originating in the bone marrow and extending to BMDMs in both sexes.

**Figure 5.**
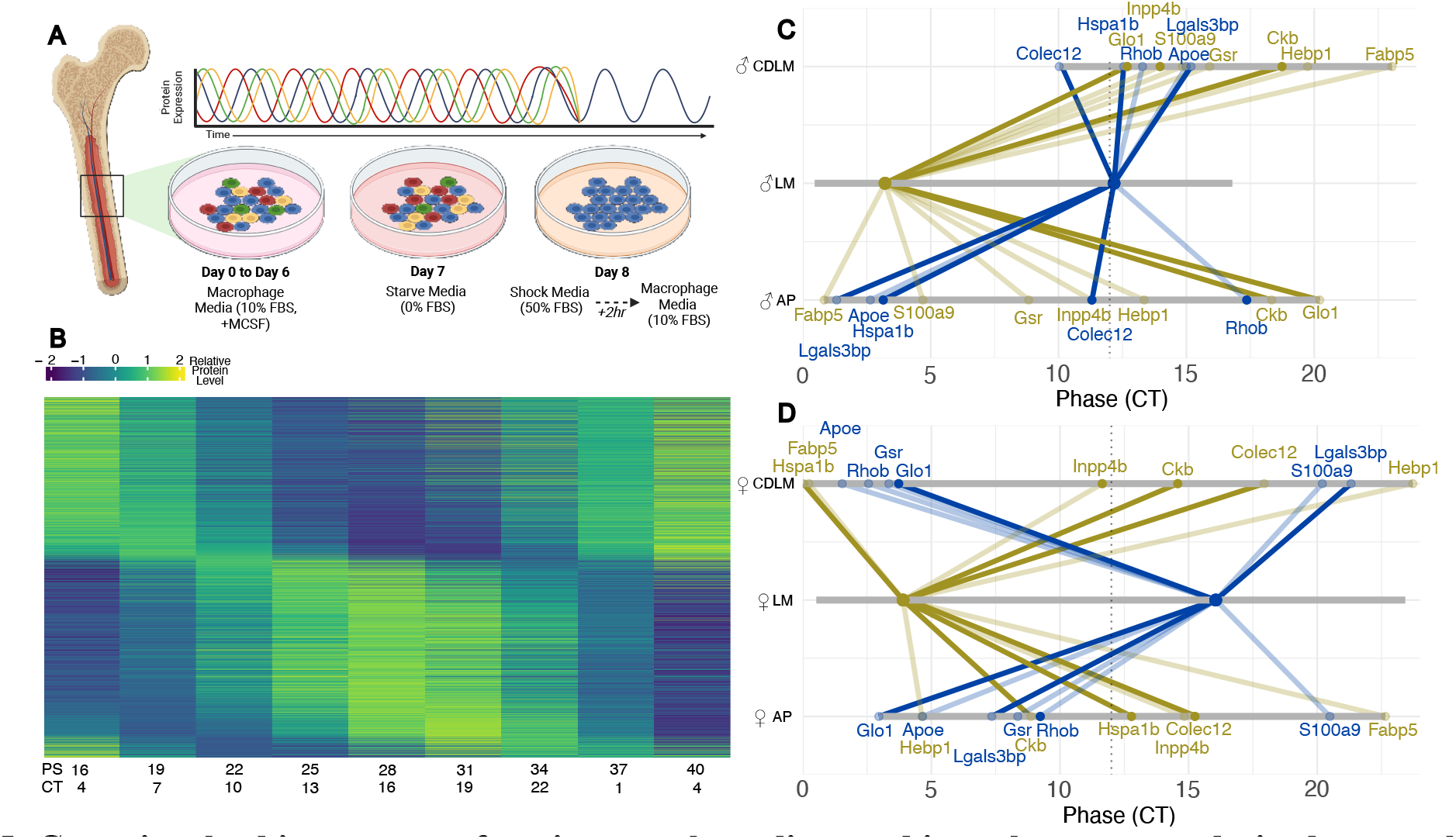
Genes involved in monocyte function are phase disrupted in male monocyte-derived macrophages. **A)** Schematic of the derivation and synchronization of bone marrow derived macrophages. Flushed bone marrow monocytes were differentiated to macrophages over a period of 6 days. On the 7th day, media with 0% FBS (starve media) was introduced. On day 8, cells were exposed to shock media supplemented with 50% FBS for two hours before returning to assay media conditions (10% FBS). **B)** Representative heatmap (PS16 through PS40) exemplifying the resultant proteomics data used for this analysis. Heatmap shown is ECHO fitted protein levels from female LM mice, sorted by peak phase. **C)** Male and **D)** Female phase change plots of macrophage proteins that were identified as DEGs in monocyte differential expression. Grey bars show range of phase, solid diagonal lines indicate circadian protein in control condition (DS/LM), lightly shaded diagonal lines indicate not circadian in DS/LM, solid dot indicates circadian in experimental condition (CD or DS/AP), lightly shaded dot indicates non-circadian in experimental condition. Blue = evening peaking DEGs, Tan = morning peaking DEGs.

## DISCUSSION

While the individual interactions between bone health, ADRDs, and circadian regulation are well-known, the concordance between these three factors is not well investigated and little is understood about the role that sex plays in this interaction. Our high-resolution μCT imaging demonstrated an effect of AP on bone deterioration in males, but not in the female AP mice (**Figure 2 A** and **C**). This divergence suggests sex-specific vulnerability to ADRD-associated skeletal deterioration. The sex difference in trabecular bone is consistent with findings in a prior study in the 5XFAD model, where 8-month-old males exhibited reduced bone health, while females showed improvement,^38^ though at 12 months, both male and female 5XFAD mice were reported to have similar degradation in bone health.^27^ Here, the around 8 month-old female mice are pre-menopause and not expected to have reduced estrogen levels, which would accelerate bone loss, a typical comorbidity in female Alzheimer’s patients.^1^ Therefore, the stress of the AP genotype in this case may not be temporally aligned with the hormonal changes that would lead to accelerated bone loss. These observations underscore the need to account for both sex and age when examining the skeletal impact of AD.

In the cortical bone, we found evidence that the bone matrix was undergoing non-enzymatic modifications, leading to overall loss of bone strength and resistance to fracture. µCT analysis revealed no structural changes in cortical morphometry, consistent with slower remodeling and bone turnover in this region^39^ (**Supplemental Figure 6**). Follow up compositional analysis using RS demonstrated that cortical bone changes were restricted to the organic matrix with the AP genotype being associated with increased accumulation of carboxymethyl-lysine (CML) (**Figure 3**). This finding, along with fAGE quantification (**Supplemental Figure 7**), points to increased collagen glycation, which can stiffen the organic matrix. Functionally, organic matrix modification translates to a measurable reduction in bone strength.^27,46^ Consequently, we confirmed that initiation and maximum toughness were significantly decreased in CD and AP groups (**Figure 3**). Decreased toughening effect, the measure of post-yield energy, was only significantly altered in the female CD and AP groups. This sex-specific CD and AP-driven reduction in extrinsic toughening highlights an increased fracture vulnerability.^47,48^ Early organic phase damage aligns with the broader metabolic and oxidative stress conditions observed in CD and AD pathology.^10^

Monocytes serve as the key intermediate in the HSC–monocyte–osteoclast differentiation axis and they had the most significant transcriptomic changes based on our scRNA-seq data (**Figure 4**). CD male monocytes showed up-regulation of inflammatory genes (*S100a8, S100a9, Ngp*), while AP male monocytes increased *Gsr* expression, which is an oxidative stress response.^19^ In the CD/AP male monocytes, we found down-regulation of *Dusp1*, a MAPK regulator that typically restricts osteoclastic bone resorption.^43^ Osteoblasts meanwhile, had too few DEGs to analyze, though their proportions were increased by AP and decreased by CD in males (**Figure 4 G**). When considered alongside the trabecular changes observed in male mice, this suggests that the inflammatory conditions in the AP male mice drive osteoclast-osteoblast activity, with resorptive activity ultimately outweighing osteoblast function. In CD male animals, preserved bone microarchitecture and reduced osteoblast proportion may reflect blunting of overall bone remodeling activity. These findings suggest that monocytes, rather than upstream progenitors, may serve as the earliest responsive cellular node to disease and environmental stress. Therefore, we were interested in the possibility of disruptions conserved in the monocyte to macrophage lineage.

Our extended study of the bone marrow to assess the circadian coordination of protein levels in monocyte-derived macrophages from both male and female mice, revealed a core set of genes that overlapped between differentially expressed monocyte transcripts and circadian macrophage proteins (**Figure 5 C** and **D**). Importantly, the time-of-day coordination of these genes (eg., *Apoe, Gsr*, and *S100a9*) were found to be strongly delayed in CD males and consistently disrupted in the AP group. Macrophage function is known to be circadianly organized, aligning immune activity with time of day.^32,45^ This implies that altered macrophage function could stem from disrupted progenitors within the bone marrow, in response to disease or environmental stress. Importantly, the macrophages in this assay were derived from animals merely *exposed* to these conditions, meaning the cells did not directly experience *in vivo* stimuli, reinforcing the concept of a lasting lineage-specific modification.

Collectively, these data indicate that the oxidative stress response detected in the bone marrow and the fAGE accumulation observed in the cortical bone likely represent shared system redox imbalance in the AP genotype. Due to scRNA-seq limitations, our analysis of bone marrow responses was restricted to males, leaving the cellular basis of female-specific adaptations unexplored at this time. It is plausible that females possess a greater cellular tolerance to oxidative stress,^49,50^ which could preserve trabecular organization despite equivalent matrix-level damage. In this framework, the maintained trabecular quality observed in females may reflect effective cellular stress responses, even as acellular matrix components accumulate damage and contribute to overall loss of bone strength.

Together, our findings in bone morphometry encompass results consistent with other models of ADRD. While the APP/PS1 mouse model may not recapitulate exact human disease, integration of results from the CD model, scRNA-seq, and circadian proteomics advances our understanding of the brain-bone axis and suggests potential mechanisms by which neurodegeneration may impact bone health. The focus on monocyte-derived macrophages as a functional unit that relies on bone resident progenitors is a crucial addition to bridging the gap in our understanding of how this disease impacts multiple systems in the body. Moreover, our approach shows that high resolution μCT, with the inclusion of scRNA-seq, are together a powerful analytical platform to understand the macro-scale health effects that arise from cell-cell interactions in the bone marrow niche. Future studies should expand on the contribution of monocyte-derived macrophages in the pathology of ADRD by exploring the temporal progression of bone-immune interactions across disease stages and sexes. Such longitudinal studies will better approximate factors in the brain-bone axis and delineate the multifaceted drivers of bone loss in ADRDs. Ultimately, this work has contributed to the foundation of understanding skeletal health in aging populations burdened by neurodegenerative and environmental stressors.

## ACKNOWLEDGEMENTS

Eduardo A.C. Almeida - Bone and Signaling Laboratory, Space Biosciences Division, NASA Ames Research Center.

## AUTHOR CONTRIBUTIONS

**Noah G. Allen:** Conceptualization; Data Curation, Formal analysis; Investigation; Methodology; Visualization; Writing - Original Draft; Writing - Review & Editing

**Carmalena V. Cordi:** Conceptualization; Data Curation, Formal analysis; Investigation; Methodology; Visualization; Writing - Review & Editing

**Joan E. Llabre:** Data Curation, Formal analysis; Investigation; Methodology; Writing - Review & Editing

**Joshua Chuah:** Data Curation

**Gretchen T. Clark:** Conceptualization; Investigation; Methodology

**Angela J. Kubik:** Conceptualization; Investigation; Methodology

**Naomi G. Falkenberg:** Investigation

**Meaghan S. Jankowski:** Conceptualization; Methodology, Resources, Writing - Review & Editing, Project administration

**Rukmani A. Cahill:** Investigation; Formal analysis; Methodology; Visualization

**Ava A. Herzog:** Investigation; Data Curation

**Mallika Subash Chander:** Data Curation; formal analysis; methodology

**Deepak Vashishth:** Conceptualization; Resources; Project administration; Writing - Review & Editing; Funding acquisition

**Jennifer M. Hurley:** Conceptualization; Resources; Writing - Review & Editing; Supervision; Project administration; Funding acquisition

**Elizabeth A. Blaber:** Conceptualization; Resources; Writing - Review & Editing; Supervision; Project administration; Funding acquisition

