## Supplemental Data for "Alzheimer’s Disease and circadian disruption sex-specifically contribute to a loss of bone maintenance in APP/PS1 model mice"

**Supplemental Table 1: 2-way ANOVA analysis and Fisher's Exact test on serum and brain Aβ42 ELISA results.** Brain homogenate data was below the assay limit of detection (LOD) resulting in zero inflation, thus analyzed via contingency test (Fisher's Exact test) to determine significant genotype effect. The LOD contingency table is shown where Aβ42 detection is reported (Aβ42 – or Aβ42 +). Mean and standard deviation reported (top), test statistic (Fstat) p-value reported (bottom). Bolded red values highlight significant 2-way ANOVA result ($p<0.05$).

| Aβ42 ELSIA | | Condition | Serum | Brain | | Aβ42 - | Aβ42 + |
|---|---|---|---|---|---|---|---|
| Mean±SD | Male | DS/LM | 62.17 ±18.31 | 0 ±0 | | 4 | 0 |
| | | DS/AP | 219.7 ±67.44 | 47.81 ±14.18 | | 0 | 4 |
| | | CD/LM | 85.36 ±7.938 | 0 ±0 | | 4 | 0 |
| | | CD/AP | 196 ±60.41 | 102.6 ±19.56 | | 0 | 4 |
| 2-way ANOVA | Lighting | Fstat | <0.001 | | Fisher's Exact (P value) | **2.00E-04** | |
| | | p.val | 0.987 | | | | |
| | Genotype | Fstat | 75.680 | | | | |
| | | p.val | **<0.0001** | | | | |
| | Interaction | Fstat | 2.315 | | | | |
| | | p.val | 0.138 | | | | |
| Mean±SD | Female | DS/LM | 90.64 ±11.8 | 0 ±0 | | 4 | 0 |
| | | DS/AP | 200.1 ±59.83 | 86.4 ±50.29 | | 0 | 4 |
| | | CD/LM | 99.37 ±18.49 | 2.03 ±4.53 | | 5 | 0 |
| | | CD/AP | 213 ±43.89 | 145.8 ±213 | | 0 | 4 |
| 2-way ANOVA | Lighting | Fstat | 0.699 | | Fisher's Exact (P value) | **<0.0001** | |
| | | p.val | 0.409 | | | | |
| | Genotype | Fstat | 74.820 | | | | |
| | | p.val | **<0.0001** | | | | |
| | Interaction | Fstat | 0.025 | | | | |
| | | p.val | 0.875 | | | | |

**Supplemental Table 2: Repeated measures, mixed effects results testing for age and treatment effects in changing activity volume.** Test statistic (Fstat) and p-value are reported. Bolded red values highlight significant mixed-effects result ($p<0.05$).

| Mixed-effects analysis | | Males | Females |
|---|---|---|---|
| Age | Fstat | 20.610 | 4.788 |
| | p.val | **0.0002** | **0.0407** |
| Treatment | Fstat | 2.584 | 0.7470 |
| | p.val | 0.0791 | 0.5357 |
| Interaction | Fstat | 0.338 | 3.107 |
| | p.val | 0.798 | **0.0496** |

**Supplemental Table 3: Mid-shaft cortical femur morphology assessed by micro-CT imaging. Results shown are mean and standard deviation for each group and parameter.** Two-way ANOVA results shown, Fstat = calculated F statistic, p.val = p-value calculated. BA/TA% = bone area fraction, Ec.Pm = endocortical perimeter, Ps.Pm = periosteal perimeter, Ct.Th = cortical thickness. Bolded red values highlight significant 2-way ANOVA result (p<0.05).

| Cortical Femur | | Condition | Ct.Ar | Ct.Ar/Tt.Ar (%) | Ct.Th | Ec.Pm | Ps.Pm | Tt.Ar |
|---|---|---|---|---|---|---|---|---|
| Mean±SD | Male | DS/LM | 1.293 ±0.186 | 61.315 ±3.162 | 0.28 ±0.029 | 3.557 ±0.211 | 5.645 ±0.326 | 2.104 ±0.238 |
| | | DS/AP | 1.289 ±0.174 | 60.723 ±3.067 | 0.275 ±0.025 | 3.693 ±0.156 | 5.734 ±0.347 | 2.119 ±0.231 |
| | | CD/LM | 1.309 ±0.211 | 60.342 ±4.727 | 0.277 ±0.036 | 3.653 ±0.223 | 5.774 ±0.345 | 2.163 ±0.235 |
| | | CD/AP | 1.383 ±0.225 | 59.93 ±3.737 | 0.29 ±0.043 | 3.65 ±0.332 | 5.875 ±0.309 | 2.238 ±0.269 |
| 2-way ANOVA | Lighting | Fstat | 0.693 | 0.486 | 0.312 | 0.112 | 1.473 | 1.197 |
| | | p.val | 0.411 | 0.491 | 0.580 | 0.740 | 0.234 | 0.282 |
| | Genotype | Fstat | 0.275 | 0.157 | 0.109 | 0.680 | 0.725 | 0.314 |
| | | p.val | 0.604 | 0.695 | 0.743 | 0.416 | 0.401 | 0.579 |
| | Interaction | Fstat | 0.344 | 0.005 | 0.702 | 0.743 | 0.003 | 0.136 |
| | | p.val | 0.562 | 0.944 | 0.408 | 0.395 | 0.959 | 0.715 |
| Mean±SD | Female | DS/LM | 1.104 ±0.045 | 64.175 ±4.268 | 0.266 ±0.03 | 2.99 ±0.16 | 4.951 ±0.167 | 1.645 ±0.121 |
| | | DS/AP | 1.049 ±0.083 | 63.242 ±1.934 | 0.262 ±0.011 | 3.042 ±0.21 | 4.944 ±0.238 | 1.66 ±0.153 |
| | | CD/LM | 1.092 ±0.076 | 65.075 ±2.484 | 0.275 ±0.002 | 2.981 ±0.187 | 4.997 ±0.202 | 1.68 ±0.129 |
| | | CD/AP | 1.117 ±0.1 | 63.585 ±2.3 | 0.263 ±0.002 | 3.019 ±0.28 | 5.101 ±0.098 | 1.756 ±0.067 |
| 2-way ANOVA | Lighting | Fstat | 0.784 | 0.310 | 0.482 | 0.035 | 1.904 | 1.822 |
| | | p.val | 0.385 | 0.583 | 0.496 | 0.853 | 0.182 | 0.191 |
| | Genotype | Fstat | 0.234 | 1.179 | 1.327 | 0.290 | 0.428 | 0.878 |
| | | p.val | 0.633 | 0.289 | 0.264 | 0.595 | 0.520 | 0.359 |
| | Interaction | Fstat | 1.532 | 0.062 | 0.353 | 0.007 | 0.567 | 0.378 |
| | | p.val | 0.229 | 0.805 | 0.559 | 0.934 | 0.459 | 0.545 |

**Supplemental Table 4:** Femoral head trabecular morphology assessed by µCT imaging. Results shown are mean and standard deviation for each group and parameter. Two-way ANOVA results shown, Fstat = calculated F statistic, p.val = p-value calculated. Conn.D. = Connectivity density, DA = Dynamic anisotropy, SMI = structural model index, Tb.N = Trabecular number, Tb.Pf = Trabecular pattern factor, Tb.Sp = Trabecular separation, Tb.Th = Trabecular thickness. Bolded red values highlight significant 2-way ANOVA result (p<0.05).

| Femoral Head | | Condition | BV/TV % | Conn.D (1/mm^3) | DA | SMI | Tb.N (1/mm) | Tb.Pf (1/mm) | Tb.Sp (mm) | Tb.Th (mm) |
|---|---|---|---|---|---|---|---|---|---|---|
| Mean±SD | Male | DS/LM | 58.388 ±7.86 | 4430.154 ±831.891 | 3.47 ±1.153 | 0.699 ±0.264 | 6.735 ±0.89 | 9.011 ±4.553 | 0.095 ±0.023 | 0.087 ±0.011 |
| | | DS/AP | 51.772 ±4.622 | 4151.766 ±490.045 | 2.469 ±0.492 | 1.057 ±0.194 | 6.257 ±0.5 | 11.325 ±1.944 | 0.091 ±0.015 | 0.083 ±0.002 |
| | | CD/LM | 58.722 ±6.951 | 5241.896 ±1345.904 | 2.956 ±0.64 | 0.744 ±0.298 | 8.24 ±2.28 | 8.507 ±3.066 | 0.074 ±0.026 | 0.074 ±0.015 |
| | | CD/AP | 51.853 ±8.464 | 4333.001 ±1200.782 | 2.127 ±0.238 | 1.297 ±0.454 | 6.74 ±1.225 | 14.731 ±6.233 | 0.081 ±0.024 | 0.077 ±0.007 |
| 2-way ANOVA | Lighting | Fstat | 0.008 | 2.144 | 2.792 | 1.730 | 4.621 | 1.011 | 4.176 | 6.895 |
| | | p.val | 0.931 | 0.153 | 0.105 | 0.198 | **0.039** | 0.323 | **0.049** | **0.013** |
| | Genotype | Fstat | 8.055 | 3.065 | 12.754 | 17.731 | 4.577 | 8.751 | 0.037 | 0.016 |
| | | p.val | **0.008** | 0.090 | **0.001** | **>0.001** | **0.040** | **0.006** | 0.849 | 0.901 |
| | Interaction | Fstat | 0.003 | 0.864 | 0.112 | 0.811 | 1.220 | 1.835 | 0.464 | 0.885 |
| | | p.val | 0.958 | 0.360 | 0.740 | 0.375 | 0.278 | 0.185 | 0.501 | 0.354 |
| Mean±SD | Female | DS/LM | 62.765 ±5.084 | 8845.434 ±1612.136 | 6.116 ±1.239 | 1.431 ±0.055 | 8.009 ±0.552 | 25.617 ±5.99 | 0.051 ±0.009 | 0.078 ±0.005 |
| | | DS/AP | 66.249 ±9.255 | 11840.062 ±4710.012 | 4.895 ±0.485 | 1.966 ±0.251 | 8.255 ±1.354 | 32.568 ±3.52 | 0.037 ±0.011 | 0.081 ±0.003 |
| | | CD/LM | 64.124 ±5.795 | 8765.438 ±1451.478 | 3.484 ±0.142 | 2.104 ±0.702 | 8.245 ±0.705 | 31.023 ±11.54 | 0.041 ±0.008 | 0.08 ±0.003 |
| | | CD/AP | 69.247 ±3.726 | 9243.726 ±2079.384 | 5.529 ±1.298 | 1.654 ±0.352 | 9.033 ±0.498 | 25.596 ±7.001 | 0.036 ±0.004 | 0.077 ±0.005 |
| 2-way ANOVA | Lighting | Fstat | 0.790 | 1.499 | 4.963 | 0.925 | 2.334 | 0.065 | 2.523 | 0.552 |
| | | p.val | 0.383 | 0.233 | **0.040** | 0.348 | 0.140 | 0.801 | 0.127 | 0.465 |
| | Genotype | Fstat | 3.085 | 2.523 | 0.845 | 0.051 | 2.429 | 0.062 | 8.312 | 0.064 |
| | | p.val | 0.092 | 0.126 | 0.371 | 0.824 | 0.133 | 0.806 | **0.009** | 0.802 |
| | Interaction | Fstat | 0.112 | 1.325 | 13.268 | 6.872 | 0.666 | 4.080 | 1.813 | 2.330 |
| | | p.val | 0.741 | 0.262 | **0.002** | **0.016** | 0.423 | 0.056 | 0.192 | 0.141 |

**Supplemental Table 5: Parameters from Raman spectroscopy and fluorescent advanced glycation end product (fAGE). Results shown are mean and standard deviation for each parameter.** Two-way ANOVA results shown, Fstat = calculated F statistic, p-value = calculated p statistic. Crystallinity reported as inv. Phosphate band, Carbonate to phosphate ratio, mineral to matrix ratio (calculated with Amide I and Amide III peaks), carboxymethyl-lysine (CML), and fluorescent advanced glycation end product (fAGE) normalized to collagen content (mg).

| Cortical Femur Molecular Parameters | | Condition | Crystallinity 1/ν1PO43 | Carbonate/ Phosphate (ν1CO32- /ν1PO43-) | Mineral/ Matrix ν2PO43- /Amide I | Mineral/ Matrix ν2PO43- /Amide III | Carboxylmethyl-lysine A1150/CH2-wag | fAGEs (ng of Quinine/mg of Collagen) |
|---|---|---|---|---|---|---|---|---|
| Mean±SD | Male | DS/LM | 0.05 ±0.00038 | 0.21 ±0.02654 | 1.48 ±0.15926 | 13.18 ±2.801 | 0.07 ±0.01659 | 70.39 ±59.34114 |
| | | DS/AP | 0.05 ±0.00059 | 0.23 ±0.01164 | 1.2 ±0.10058 | 10.63 ±2.68171 | 0.14 ±0.04505 | 466.38 ±337.75185 |
| | | CD/LM | 0.05 ±0.00017 | 0.2 ±0.02212 | 1.04 ±0.28179 | 11.55 ±1.42442 | 0.12 ±0.03187 | 369.48 ±197.33318 |
| | | CD/AP | 0.05 ±0.00076 | 0.23 ±0.06318 | 1.29 ±0.14583 | 10.49 ±2.20634 | 0.15 ±0.05064 | 363.27 ±159.68415 |
| 2-way ANOVA | Lighting | Fstat | 0.123 | 0.005 | 0.532 | 4.028 | 2.205 | 0.844 |
| | | p.val | 0.732 | 0.946 | 0.480 | 0.068 | 0.163 | 0.376 |
| | Genotype | Fstat | 0.011 | 1.301 | 2.208 | 0.048 | 7.029 | 3.338 |
| | | p.val | 0.917 | 0.276 | 0.163 | 0.830 | **0.021** | 0.093 |
| | Interaction | Fstat | 0.183 | 0.110 | 0.371 | 9.042 | 0.779 | 3.555 |
| | | p.val | 0.677 | 0.746 | 0.554 | **0.011** | 0.395 | 0.084 |
| Mean±SD | Female | DS/LM | 0.053 ±0.0004 | 0.221 ±0.0264 | 12.666 ±2.5792 | 1.23 ±0.2229 | 0.052 ±0.026 | 30.626 ±17.4791 |
| | | DS/AP | 0.053 ±0.0004 | 0.233 ±0.0767 | 9.576 ±1.0483 | 0.968 ±0.344 | 0.12 ±0.0333 | 138.007 ±61.9243 |
| | | CD/LM | 0.053 ±0.0008 | 0.21 ±0.0727 | 9.8 ±1.8036 | 0.963 ±0.1976 | 0.066 ±0.0385 | 91.668 ±84.6336 |
| | | CD/AP | 0.053 ±0.0004 | 0.26 ±0.0507 | 9.93 ±2.2925 | 1.044 ±0.1575 | 0.149 ±0.0346 | 329.454 ±222.3762 |
| 2-way ANOVA | Lighting | Fstat | 0.045 | 0.086 | 1.941 | 0.788 | 2.045 | 5.320 |
| | | p.val | 0.834 | 0.773 | 0.183 | 0.388 | 0.172 | **0.037** |
| | Genotype | Fstat | 0.499 | 1.332 | 2.695 | 0.707 | 25.094 | 9.942 |
| | | p.val | 0.490 | 0.265 | 0.120 | 0.413 | **1.28E-04** | **0.007** |
| | Interaction | Fstat | 0.015 | 0.515 | 3.186 | 2.523 | 0.238 | 1.419 |
| | | p.val | 0.905 | 0.484 | 0.093 | 0.132 | 0.632 | 0.253 |

**Supplemental Table 6: Notched three point bending strength test results.** Data shown are mean and standard deviation for each group and parameter. Two-way ANOVA results shown, Fstat = calculated F statistic, p.val = p-value calculated. Bolded red values highlight significant 2-way ANOVA result ($p<0.05$).

| Femur Strength Testing | | Condition | Initiation Toughness Kc, in (MPa√m) | Maximum Toughness Kc, max (MPa√m) | Toughening Effect ΔK (MPa√m) |
|---|---|---|---|---|---|
| Mean±SD | Male | LM | 0.74 ±0.09995 | 1.76 ±0.29265 | 1.02 ±0.29116 |
| | | AP | 0.33 ±0.10815 | 0.94 ±0.44614 | 0.61 ±0.35975 |
| | | CD | 0.23 ±0.04172 | 0.92 ±0.26741 | 0.69 ±0.24902 |
| | | CD/AP | 0.18 ±0.05994 | 0.69 ±0.2113 | 0.51 ±0.18336 |
| 2-way ANOVA | Lighting | Fstat | 69.964 | 13.100 | 2.686 |
| | | p.val | **1.37E-06** | **0.003** | 0.125 |
| | Genotype | Fstat | 34.650 | 12.161 | 4.942 |
| | | p.val | **5.35E-05** | **0.004** | **0.045** |
| | Interaction | Fstat | 21.341 | 3.826 | 0.733 |
| | | p.val | **4.80E-04** | 0.072 | 0.407 |
| Mean±SD | Female | LM | 0.608 ±0.2093 | 1.811 ±0.4273 | 1.203 ±0.2951 |
| | | AP | 0.368 ±0.0902 | 1.163 ±0.3588 | 0.795 ±0.2794 |
| | | CD | 0.264 ±0.0477 | 0.723 ±0.1236 | 0.459 ±0.084 |
| | | CD/AP | 0.231 ±0.0901 | 0.73 ±0.2906 | 0.499 ±0.2078 |
| 2-way ANOVA | Lighting | Fstat | 19.815 | 27.899 | 24.621 |
| | | p.val | **4.66E-04** | **9.22E-05** | **1.71E-04** |
| | Genotype | Fstat | 6.403 | 4.960 | 3.081 |
| | | p.val | **0.023** | **0.042** | 0.100 |
| | Interaction | Fstat | 3.666 | 5.183 | 4.583 |
| | | p.val | 0.075 | **0.038** | **0.049** |

**Supplemental Table 7: Significant gene ontology (GO) terms for genes identified in the LRT analysis of aggregated bone marrow expression.** Gene sets were submitted to GO analysis in clusters based on the hierarchical expression clustering as presented in Figure 3. Term: GO term name, Source: origin of GO term (BP: Biological process, CC: Cellular Component, MF: Molecular Function), Adjusted p-value: default method implemented by gprofiler2 package.

| Cluster | Term | Source | Adjusted p-value | Gene Ratio |
|---|---|---|---|---|
| 1 | regulation of release of sequestered calcium ion into cytosol | GO:BP | 2.56E-02 | 2/4 |
| 3 | paraspeckles | GO:CC | 1.71E-03 | 2/13 |
| 3 | identical protein binding | GO:MF | 1.37E-02 | 7/13 |
| 3 | protein binding | GO:MF | 4.42E-02 | 12/13 |
| 6 | response to bacterium | GO:BP | 1.62E-05 | 9/20 |
| 6 | response to biotic stimulus | GO:BP | 1.87E-05 | 11/20 |
| 6 | calprotectin complex | GO:CC | 8.53E-05 | 2/21 |
| 6 | calcium-dependent protein binding | GO:MF | 2.21E-04 | 4/21 |
| 6 | response to other organism | GO:BP | 2.32E-04 | 10/20 |
| 6 | response to external biotic stimulus | GO:BP | 2.37E-04 | 10/20 |
| 6 | antimicrobial humoral response | GO:BP | 2.93E-04 | 5/20 |
| 6 | innate immune response | GO:BP | 3.87E-04 | 8/20 |
| 6 | biological process involved in interspecies interaction between organisms | GO:BP | 5.15E-04 | 10/20 |
| 6 | extracellular space | GO:CC | 6.23E-04 | 9/21 |
| 6 | response to external stimulus | GO:BP | 7.57E-04 | 11/20 |
| 6 | response to lipopolysaccharide | GO:BP | 9.74E-04 | 6/20 |
| 6 | response to molecule of bacterial origin | GO:BP | 1.19E-03 | 6/20 |
| 6 | cytoplasm | GO:CC | 1.20E-03 | 19/21 |
| 6 | defense response to symbiont | GO:BP | 1.32E-03 | 8/20 |
| 6 | ion binding | GO:MF | 1.78E-03 | 15/21 |
| 6 | regulation of transport | GO:BP | 2.41E-03 | 9/20 |
| 6 | antimicrobial humoral immune response mediated by antimicrobial peptide | GO:BP | 2.79E-03 | 4/20 |
| 6 | small molecule binding | GO:MF | 2.90E-03 | 15/21 |
| 6 | defense response to other organism | GO:BP | 2.95E-03 | 8/20 |
| 6 | humoral immune response | GO:BP | 3.08E-03 | 5/20 |
| 6 | defense response | GO:BP | 3.74E-03 | 9/20 |
| 6 | regulation of vesicle-mediated transport | GO:BP | 4.39E-03 | 6/20 |
| 6 | calcium ion binding | GO:MF | 5.59E-03 | 6/21 |
| 6 | response to stress | GO:BP | 5.70E-03 | 12/20 |
| 6 | neutrophil aggregation | GO:BP | 7.46E-03 | 2/20 |
| 6 | specific granule | GO:CC | 7.72E-03 | 2/21 |
| 6 | extracellular region | GO:CC | 9.27E-03 | 9/21 |
| 6 | regulation of inflammatory response | GO:BP | 1.09E-02 | 5/20 |

| | | | | |
|---|---|---|---|---|
| 6 | cytolysis by host of symbiont cells | GO:BP | 1.12E-02 | 2/20 |
| 6 | defense response to bacterium | GO:BP | 1.17E-02 | 5/20 |
| 6 | programmed cell death | GO:BP | 1.28E-02 | 9/20 |
| 6 | cell death | GO:BP | 1.28E-02 | 9/20 |
| 6 | regulation of response to external stimulus | GO:BP | 1.36E-02 | 7/20 |
| 6 | regulation of localization | GO:BP | 1.37E-02 | 9/20 |
| 6 | antioxidant activity | GO:MF | 1.45E-02 | 3/21 |
| 6 | cytoplasmic vesicle | GO:CC | 1.55E-02 | 8/21 |
| 6 | intracellular vesicle | GO:CC | 1.58E-02 | 8/21 |
| 6 | response to lipid | GO:BP | 1.79E-02 | 7/20 |
| 6 | autocrine signaling | GO:BP | 2.09E-02 | 2/20 |
| 6 | vesicle | GO:CC | 2.54E-02 | 8/21 |
| 6 | positive regulation of cellular component organization | GO:BP | 2.61E-02 | 7/20 |
| 6 | positive regulation of intrinsic apoptotic signaling pathway | GO:BP | 2.64E-02 | 3/20 |
| 6 | regulation of defense response | GO:BP | 3.05E-02 | 6/20 |
| 6 | inflammatory response | GO:BP | 3.47E-02 | 6/20 |
| 6 | positive regulation of multicellular organismal process | GO:BP | 3.69E-02 | 8/20 |
| 6 | regulation of multicellular organismal process | GO:BP | 3.99E-02 | 10/20 |
| 6 | response to toxic substance | GO:BP | 4.66E-02 | 4/20 |

**Supplemental Table 8: Pairwise testing results of antioxidant genes comparing the experimental group to control group.** Log2FC and Adjusted p-values are calculated using the FindMarkers (Seurat, v5) implementation of the MAST algorithm. Bolded red values indicate significant result (1.3<|Log2FC|) and adjusted p value <0.05). Grey boxes indicate comparisons not calculated due to low transcript abundance or pre-filtering of Log2FC(|Log2FC| < 0.1).

| Antioxidant Genes | Gene Sub-group annotation | APvControl | | CDvControl | | CD/APvControl | |
|---|---|---|---|---|---|---|---|
| | | Log2FC | Adjusted p value | Log2FC | Adjusted p value | Log2FC | Adjusted p value |
| ***Als2*** | ROS Stress Response | 0.12 | 1.00 | 0.16 | 1.00 | -0.30 | 1.00 |
| ***Apc*** | Peroxidase | | | -0.12 | 1.00 | -0.27 | 1.00 |
| ***Apoe*** | ROS Stress Response | -1.23 | 1.00 | -0.49 | 1.00 | **-2.23** | **8.34E-17** |
| ***Ctsb*** | Peroxidase | -0.20 | 1.00 | | | -0.20 | 1.00 |
| ***Cyba*** | NADPH Oxidase | | | | | 0.16 | 0.86 |
| ***Ehd2*** | Peroxiredoxin | -0.17 | 1.00 | | | -0.85 | 1.00 |
| ***Gpx1*** | Glutathione | -0.38 | 2.22458E -24 | 0.12 | 5.95E-11 | | |
| ***Gpx4*** | Glutathione | | | -0.12 | 1.00 | 0.17 | 1.00 |
| ***Gsr*** | Glutathione | **2.98** | **3.68E-286** | 0.21 | 1.00 | **1.97** | **8.71E-133** |
| ***Kif9*** | Peroxidase | 0.63 | 1.00 | -0.24 | 1.00 | -0.12 | 1.00 |
| ***Mpo*** | Peroxidase | **-1.51** | **1.87E-08** | -0.29 | 1.00 | -0.81 | 1.00 |
| ***Ncf2*** | NADPH Oxidase | 0.15 | 1.00 | 0.21 | 3.06E-11 | -0.21 | 1.00 |
| ***Nox1*** | NADPH Oxidase | 0.43 | 1.00 | 0.14 | 1.00 | 0.19 | 1.00 |
| ***Nxn*** | Thioredoxin | 0.22 | 1.00 | 0.26 | 1.00 | -0.17 | 1.00 |
| ***Prdx1*** | Peroxiredoxin | -0.42 | 2.21E-03 | -0.23 | 1.00 | 0.19 | 1.00 |
| ***Prdx2*** | Peroxiredoxin | -0.46 | 1.00 | | | 0.39 | 0.80 |
| ***Prdx3*** | Peroxiredoxin | -0.27 | 1.00 | | | | |
| ***Prdx6*** | Peroxiredoxin | | | 0.17 | 1.00 | -0.23 | 1.00 |
| ***Ptgs2*** | Peroxiredoxin | -0.27 | 1.00 | | | -0.72 | 1.00 |
| ***Rag2*** | Peroxiredoxin | 0.19 | 1.00 | -0.21 | 1.00 | 0.72 | 1.00 |
| ***Scd1*** | Metabolic Redox Enzyme | -0.85 | 0.01 | -0.13 | 1.00 | -0.64 | 0.13 |
| ***Serpinb1a*** | Antioxidant | | | 0.42 | 8.02E-05 | 0.12 | 1.00 |
| ***Slc41a3*** | Peroxidase | -0.22 | 1.00 | 0.22 | 1.00 | 0.11 | 1.00 |
| ***Sod1*** | Superoxide Dismutase | | | | | -0.15 | 1.00 |
| ***Txnip*** | ROS Stress Response | | | | | 0.17 | 1.00 |

**Supplemental Table 9: Significant differentially expressed genes (DEGs, as compared to control group) for each condition and the overlap between DEGs across the conditions.** Enriched Gene Ontology (GO) for biological processes shown for unique gene expression sets. Intersection Set: transcripts contributing to GO terms. Adjusted p-value: default method implemented by clusterProfiler package.

| Intersection Group | Transcripts | GO ID | Description | Intersection Set | Adjusted p-value | Gene Ratio |
|---|---|---|---|---|---|---|
| Shared across all | *Hspa1b, Hspa1a, Egr1* | | | | | |
| Unique to AP | *Gsr, Colec12, Dhx40, Nme7, Wrn, AA465934, Gm12166, Inpp4b, Slc15a2, Cdk5rap1, Chil3, Zfp985, Gm18870, Gm4070, Hpgd, Lgals3bp, Gm42031* | GO:0005044 | scavenger receptor activity | *Lgals3bp/Colec12* | 0.0036 | 2/13 |
| | | GO:0008408 | 3'-5' exonuclease activity | *Nme7/Wrn* | 0.0096 | 2/13 |
| | | GO:0038024 | cargo receptor activity | *Lgals3bp/Colec12* | 0.0107 | 2/13 |
| | | GO:0004527 | exonuclease activity | *Nme7/Wrn* | 0.0107 | 2/13 |
| | | GO:0004386 | helicase activity | *Dhx40/Wrn* | 0.0243 | 2/13 |
| Unique to CD/AP | *Pde4d, Gm11361, Dusp1, Apoe, Fabp5, Gm26532, Rgcc, Ppfia4, Apoc2, Ppp1r15a, Sirpb1c, Hmga1b, Adssl1, Glo1, Rnase2a, Gm50022, Gm10503, Ccdc180, Pmaip1, Btla, Ccl2, Adgrg5, Gm36560, Ckb, Slc20a1, Rwdd2a, Coq10b, Tppp3, H2-DMb2, Smim8, Ets2, Mafb, Trim36, Trib1* | GO:0001933 | negative regulation of protein phosphorylation | *Pde4d/Apoe/Dusp1/Ppp1r15a/Trib1* | 0.0102 | 5/29 |
| | | GO:0042326 | negative regulation of phosphorylation | *Pde4d/Apoe/Dusp1/Ppp1r15a/Trib1* | 0.0102 | 5/29 |
| | | GO:0071830 | triglyceride-rich lipoprotein particle clearance | *Apoe/Apoc2* | 0.0102 | 2/29 |
| | | GO:0034384 | high-density lipoprotein particle clearance | *Apoe/Apoc2* | 0.0102 | 2/29 |
| | | GO:0010563 | negative regulation of phosphorus metabolic process | *Pde4d/Apoe/Dusp1/Ppp1r15a/Trib1* | 0.0102 | 5/29 |
| Unique to CD | *Hbegf, S100a8, S100a9, Vcan, Ngp, Mmp8, B930036N10Rik, Lcn2, Wfdc21, Hist1h2bc, Hist1h2ac, Gm13814, Hist1h2br, Hist1h3d, Gm11290, Hist2h4, Hist1h2bg, Vill, Gm26887, D830025C05Rik, Hist1h2bn, Camp, Hist2h2be, Hist1h4n, Hist1h2ah, Gm37168, Gm49692, Frmd8os* | GO:0018119 | peptidyl-cysteine S-nitrosylation | *S100a8/S100a9* | 0.0022 | 2/14 |
| | | GO:0017014 | protein nitrosylation | *S100a8/S100a9* | 0.0022 | 2/14 |
| | | GO:0032496 | response to lipopolysaccharide | *S100a8/S100a9/Mmp8/Wfdc21* | 0.0022 | 4/14 |
| | | GO:0070486 | leukocyte aggregation | *S100a8/S100a9* | 0.0022 | 2/14 |
| | | GO:0002237 | response to molecule of bacterial origin | *S100a8/S100a9/Mmp8/Wfdc21* | 0.0022 | 4/14 |
| AP & CD | *Jun, Rhob, Gm49980, Gm43305, Tex14* | | | | | |
| CD/AP & AP | *Gm10260, Emp1, Hebp1, Gm42047, Nrxn1* | | | | | |
| CD/AP & CD | *Klf2, Hist1h4m, Tmem108* | | | | | |

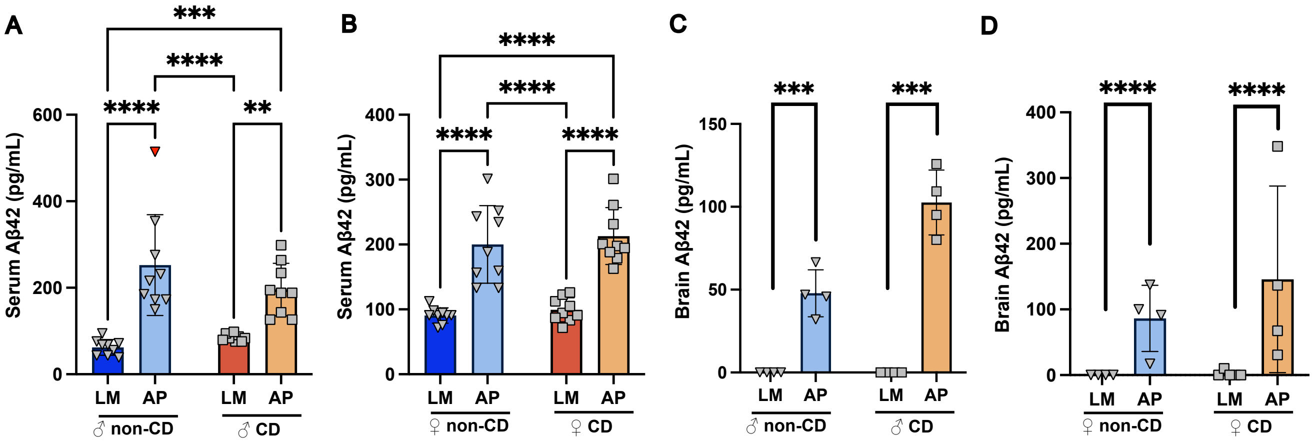

**Supplemental Figure 1: APP/PS1 mouse (AP) show significant increase of amyloid beta 42 in the circulation and brain tissue.** Amyloid beta 42 quantified via ELISA in **A)** male serum, **B)** female serum, Tukey's post-hoc pairwise comparison displayed (* p<0.05, **p<0.01, ***p<0.001, ****p<0.0001). **C)** male brain homogenate, **D)** female brain homogenate. In LM brain homogenate, values were below assay limit of detection, significance indicates results from Fisher's exact test based on frequency (detected vs. not detected) (* p<0.05, **p<0.01, ***p<0.001, ****p<0.0001). Dark blue = control/littermates (LM), Light blue = Alzheimer's genotype (AP), Dark orange = Circadianly disrupted (CD), and Light orange = CD/AP.

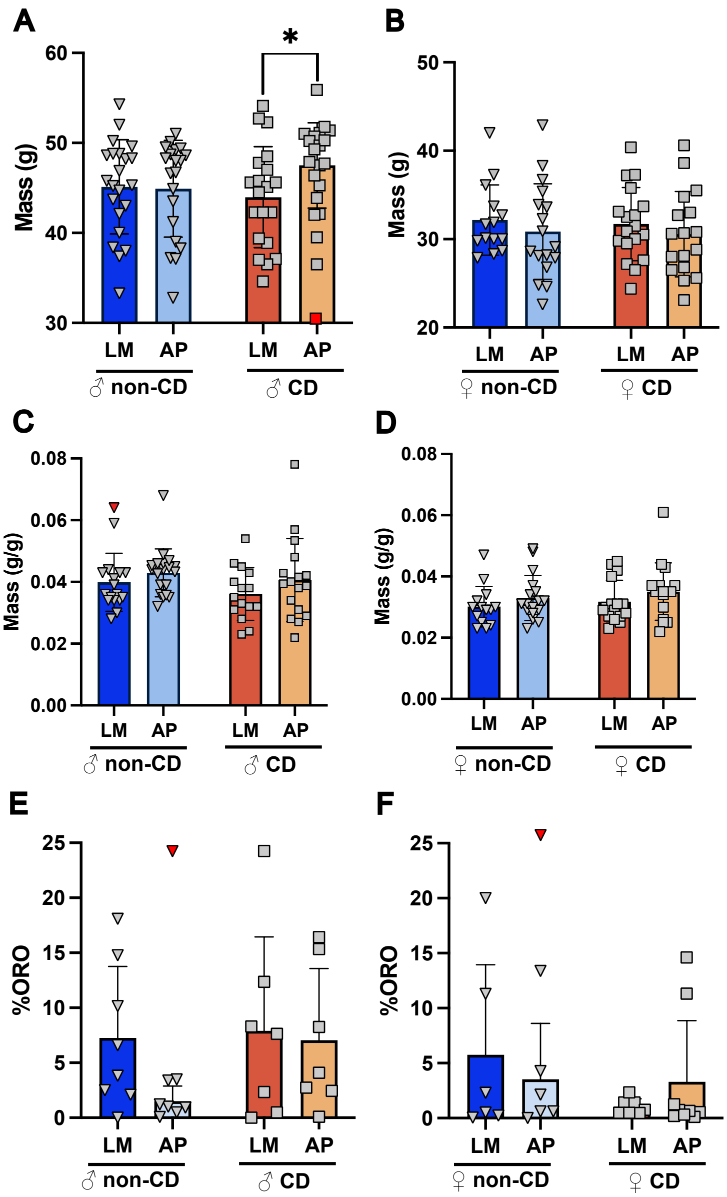

**Supplemental Figure 2: Body mass, liver mass, and liver lipids show no signs of metabolic effects due to CD and AP.** **A)** Male and **B)** Female body mass at the end of the 22-week protocol. **C)** Male and **D)** female liver mass normalized to body mass at the end of the 22-week protocol. **E)** Male and **F)** female quantification of liver lipids by cryosectioning and staining with Oil Red O. Tukey's post-hoc pairwise comparison displayed (* $p<0.05$), Red points indicate outlier removed from analysis as determined by Grubb's test. Dark blue = control/littermates (LM), Light blue = Alzheimer's genotype (AP), Dark orange = Circadianly disrupted (CD), and Light orange = CD/AP.

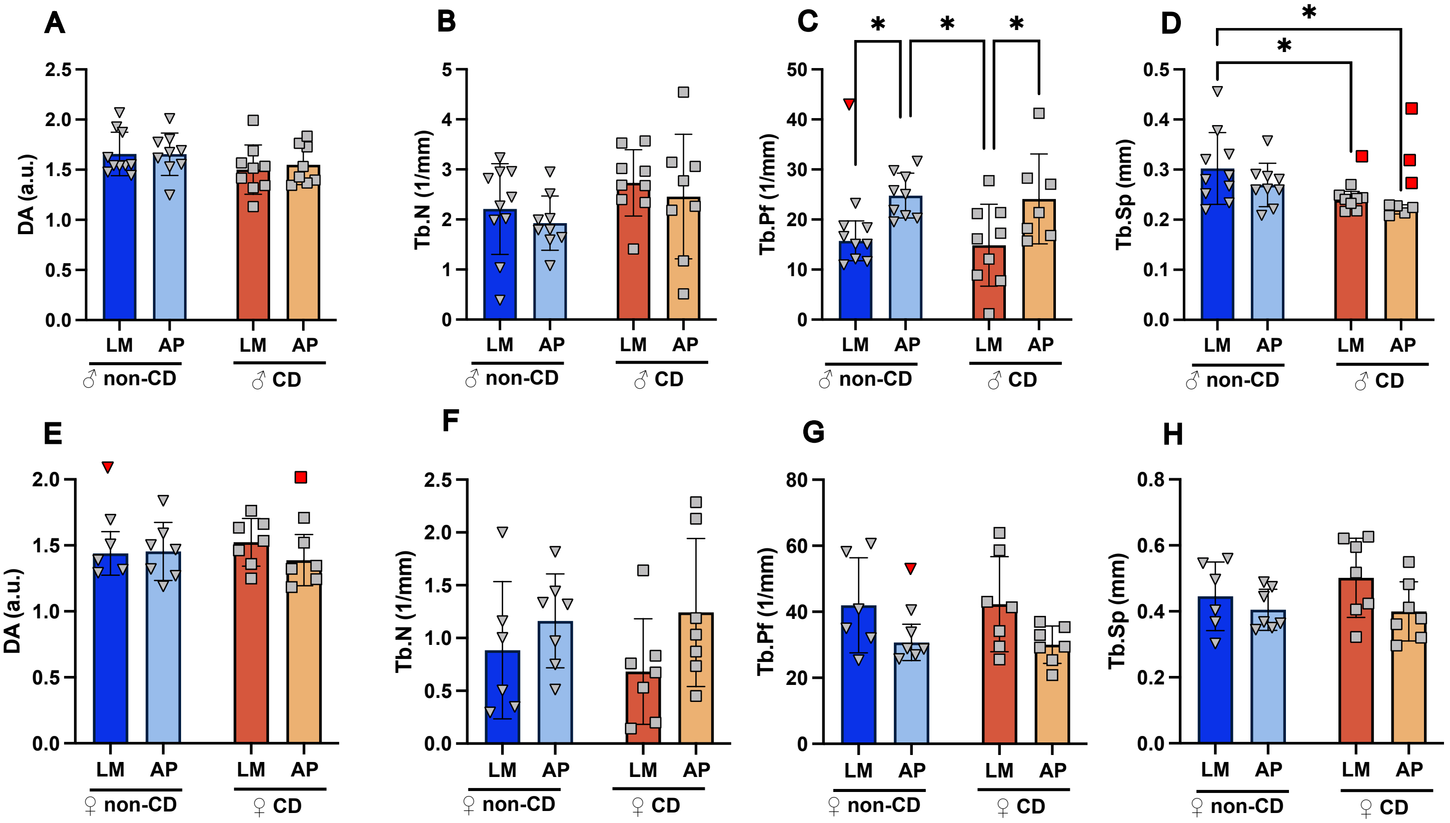

**Supplemental Figure 3: Distal femur µCT parameters are sex-specifically affected by AP and CD.** Male A) DA = Dynamic Anisotropy, B) Tb.N = Trabecular Number, C) Tb.Pf = Trabecular Pattern Factor, and D) Tb.Sp = Trabecular Separation by genotype and lighting condition. Female E) DA, F) Tb.N, G) Tb.Pf, and H) Tb.Sp by genotype and lighting condition. Tukey's post-hoc pairwise comparison displayed (* $p<0.05$). Red points indicate outlier removed from analysis as determined by Grubb's test. Dark blue = control/littermates (LM), Light blue = Alzheimer's genotype (AP), Dark orange = Circadianly disrupted (CD), and Light orange = CD/AP.

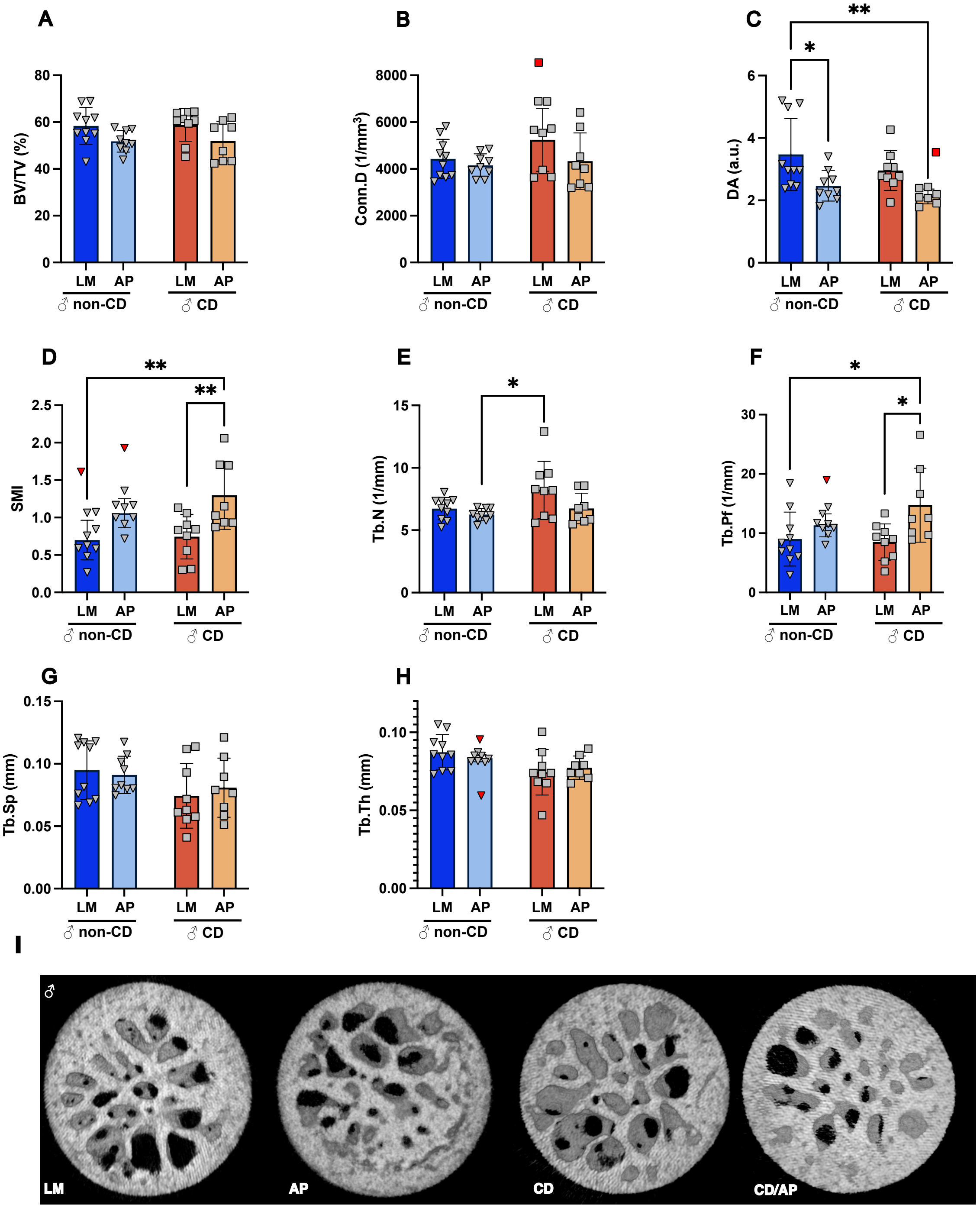

**Supplemental Figure 4: Male femoral head trabecular morphology is affected by AP**. The A) Bone volume fraction (BV/TV%), B) Connectivity density (Conn.D), C) Degree of Anisotropy (DA), D) Structural model index (SMI), E) Trabecular Number (Tb.N), F Trabecular Pattern Factor (Tb.Pf), G) Trabecular Separation (Tb.Sp), and H) Trabecular Thickness (Tb.Th) of the male femoral head trabeculae. I) Representative reconstruction of volume of interest in femoral head from male animals. Tukey's post-hoc pairwise comparison displayed (* $p<0.05$, ** $p<0.01$). Red points indicate outlier removed from analysis as determined by Grubb's test. Dark blue = control/littermates (LM), Light blue = Alzheimer's genotype (AP), Dark orange = Circadianly disrupted (CD), and Light orange = CD/AP.

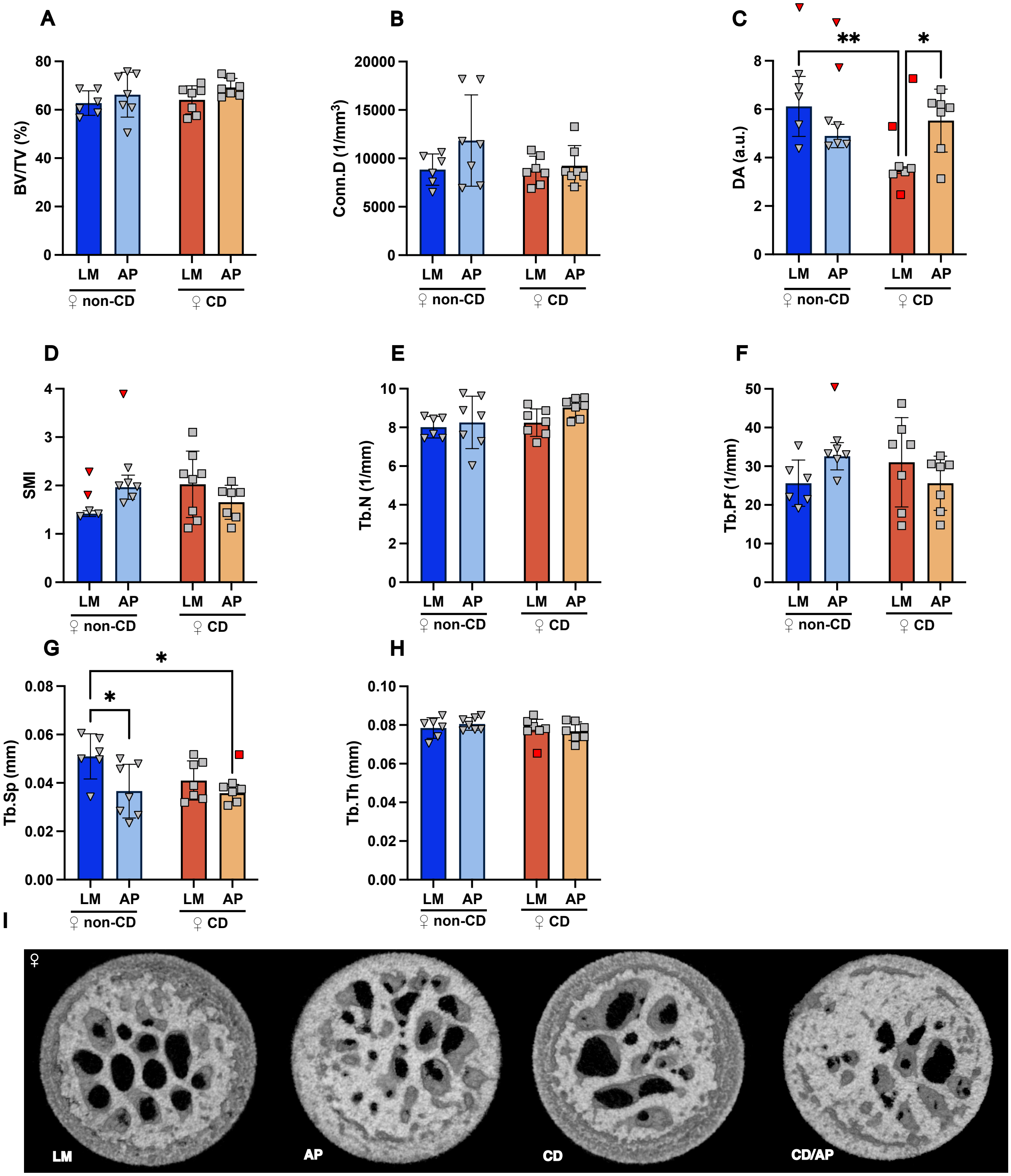

**Supplemental Figure 5: Female femoral head trabecular morphology is affected by AP**. The A) Bone volume fraction (BV/TV%), B) Connective density (Conn.D), C) Degree of Anisotropy (DA), D) Structural model index (SMI), E) Trabecular Number (Tb.N), F Trabecular Pattern Factor (Tb.Pf), G) Trabecular Separation (Tb.Sp), and H) Trabecular Thickness (Tb.Th) of the female femoral head trabeculae. I) Representative reconstruction of volume of interest in femoral head from female animals. Tukey's post-hoc pairwise comparison displayed (* p<0.05, **p<0.01). Red points indicate outlier removed from analysis as determined by Grubb's test. Dark blue = control/littermates (LM), Light blue = Alzheimer's genotype (AP), Dark orange = Circadianly disrupted (CD), and Light orange = CD/AP.

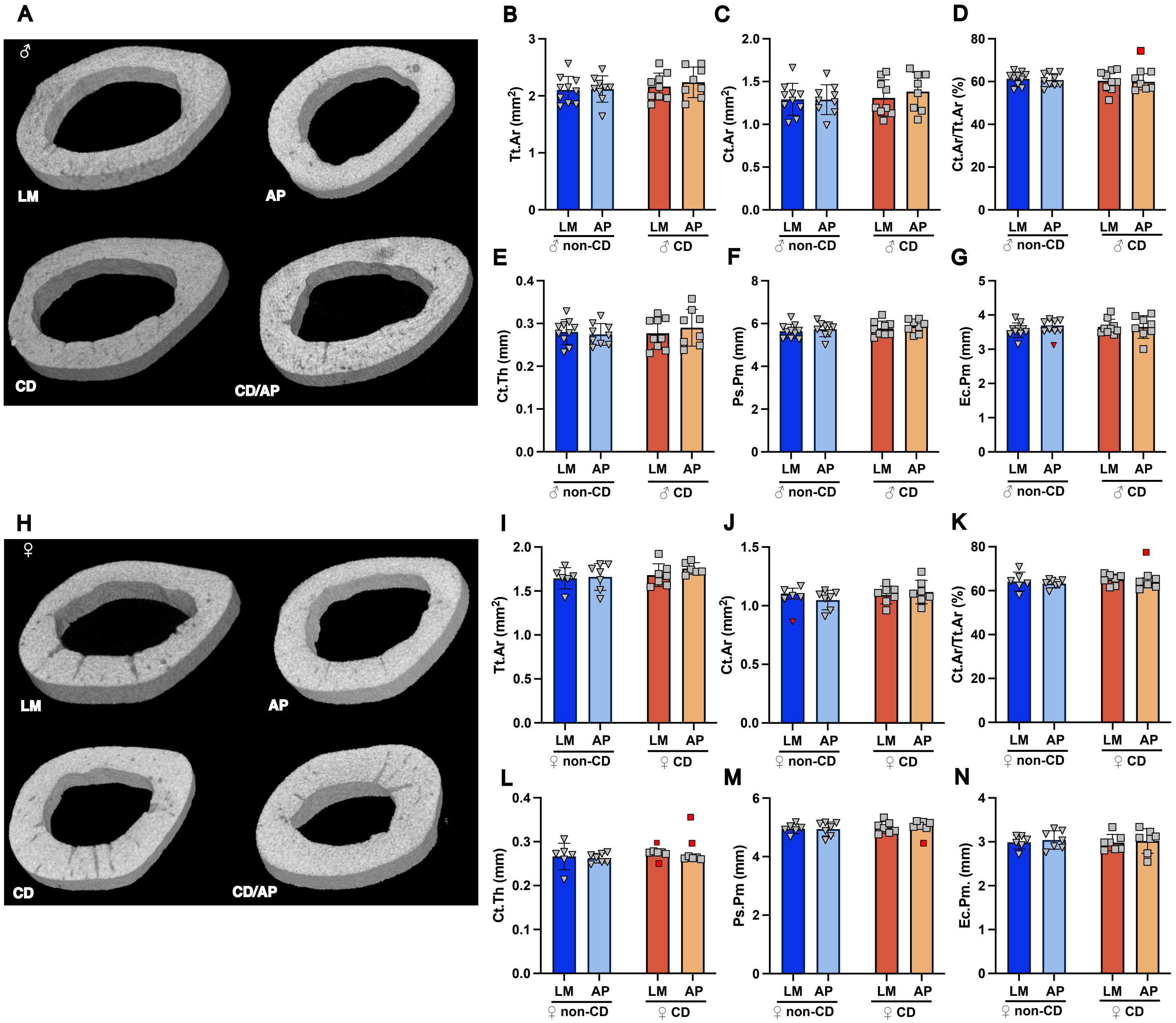

**Supplemental Figure 6: Mid-shaft cortical bone shows no effect of AP and CD.** Cortical bone morphology in male mice: A) Example reconstruction for male cortical volume of interest, B) Total cross sectional area inside the periosteal envelope (Tt.Ar), C) Cortical bone area (Ct.Ar), D) Cortical area fraction (Ct.Ar/Tt.Ar) E) Cortical thickness (Ct.Th) F) Periosteal Perimeter (Ps.Pm). G) Endocortical perimeter (Ec.Pm). Cortical bone morphology in female mice: H) Example reconstruction for female cortical volume of interest, I) Total cross sectional area inside the periosteal envelope (Tt.Ar), J) Cortical bone area (Ct.Ar), K) Cortical area fraction (Ct.Ar/Tt.Ar) L) Cortical thickness (Ct.Th) M) Periosteal Perimeter (Ps.Pm). N) Endocortical perimeter (Ec.Pm). Tukey's post-hoc pairwise comparison displayed (* $p<0.05$). Red points indicate outlier removed from analysis as determined by Grubb's test. Dark blue = control/littermates (LM), Light blue = Alzheimer's genotype (AP), Dark orange = Circadianly disrupted (CD), and Light orange = CD/AP.

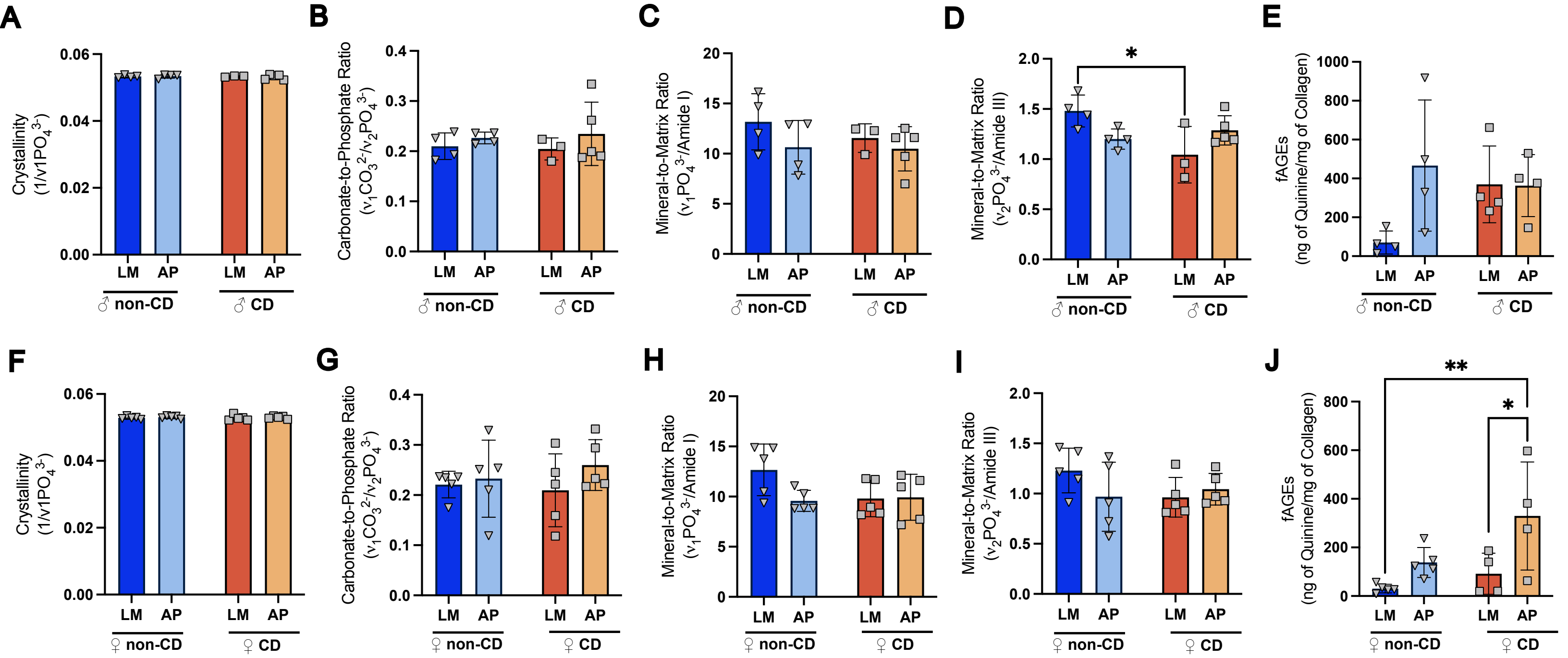

**Supplemental Figure 7: Genotype and lighting condition have a sex-specific effect on fAGEs in cortical bone.** The results of Raman spectroscopy and fluorescent AGE quantification. A) crystallinity, B) carbonate to phosphate ratio, C) mineral to matrix ratio-Amide 1, D) mineral to matrix ratio-Amide 3, and E) fAGE levels for male bones. F) crystallinity, G) carbonate to phosphate ratio, H) mineral to matrix ratio-Amide 1, I) mineral to matrix ratio-Amide 3, and J) fAGE levels for female bones. Dark blue = control/littermates (LM), Light blue = Alzheimer's genotype (AP), Dark orange = Circadianly disrupted (CD), and Light orange = CD/AP.

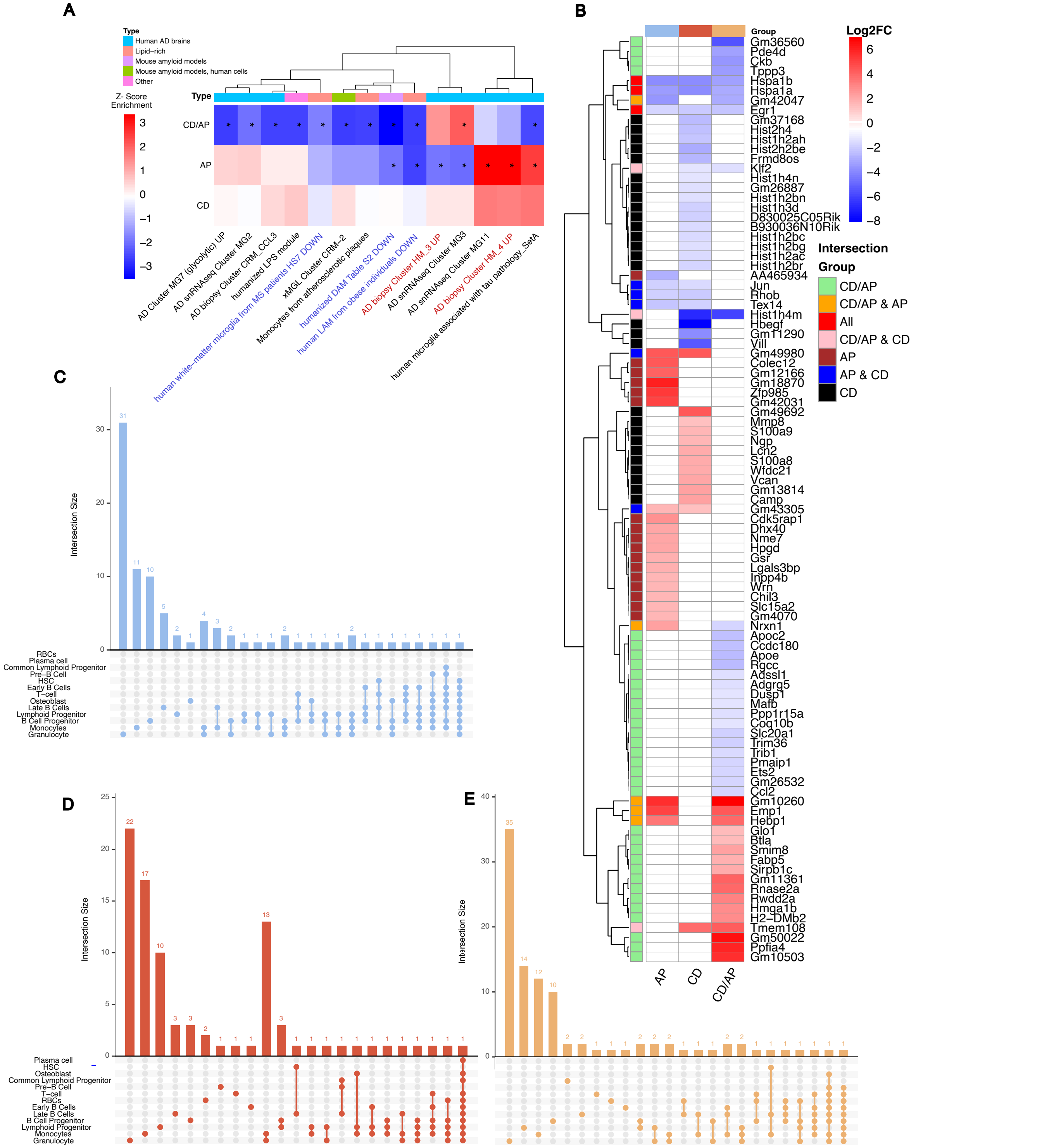

**Supplemental Figure 8: DEGs are cell type specific to AP and CD; AP monocytes are enriched for Alzheimer's transcriptomic programs. A)** GSEA enrichment using the MAST implementation, comparing monocyte enrichment to Alzheimer's patient transcriptomic gene sets derived from Goate et. al, **B)** Monocyte DEG Log2FC, white indicates no significant change. **C)** The overlap DEGs between cell types visualized with each cell type plotted with UpsetR (v1.4.0) Light blue = Alzheimer's genotype (AP) VS LM, **D)** Dark orange = Circadianly disrupted (CD) vs LM, and **E)** Light orange = CD/AP vs LM.

**Supplemental Datasets**

Marker genes used to identify cell clusters, cross referenced with CellxGene database.

| Cluster | Gene | Name |
|---|---|---|
| B Cell Progenitor | Bach2 | BTB and CNC homology, basic leucine zipper transcription factor 2 |
| | Vpreb1 | V-set pre-B cell surrogate light chain 1 |
| | Lef1 | lymphoid enhancer binding factor 1 |
| | Cecr2 | CECR2, histone acetyl-lysine reader |
| | Vpreb3 | V-set pre-B cell surrogate light chain 3 |
| CLP | Igll1 | immunoglobulin lambda-like polypeptide 1 |
| | Lef1 | lymphoid enhancer binding factor 1 |
| | Dntt | deoxynucleotidyltransferase, terminal |
| | Gfra2 | glial cell line derived neurotrophic factor family receptor alpha 2 |
| Early B Cells | Ms4a1 | Membrane Spanning 4-Domains A1 |
| | Cd79a | CD79A antigen (immunoglobulin-associated alpha) |
| | Bank1 | B cell scaffold protein with ankyrin repeats 1 |
| | Ly6d | lymphocyte antigen 6 family member D |
| Granulocyte | Lta4h | leukotriene A4 hydrolase |
| | Chil3 | chitinase-like 3 |
| | Camp | Cathelicidin Antimicrobial Peptide |
| | Ltf | lactotransferrin |
| HSC | Cd34 | CD34 antigen |
| | Gata2 | GATA binding protein 2 |
| Late B Cells | H2-Eb1 | histocompatibility 2, class II antigen E beta |
| | H2-Aa | histocompatibility 2, class II antigen A, alpha |
| Lymphoid Progenitor | Top2a | topoisomerase (DNA) II alpha |
| | Bcl7a | B cell CLL/lymphoma 7A |
| | Il7r | interleukin 7 receptor |
| | Pclaf | PCNA clamp associated factor |
| | Ezh2 | enhancer of zeste 2 polycomb repressive complex 2 subunit |
| HSC | Cd34 | CD34 antigen |
| | Gata2 | GATA binding protein 2 |
| Macrophage/ Dendritic Cells | Itgam | integrin alpha M |
| | Anpep | alanyl aminopeptidase, membrane |
| | Cd33 | CD33 molecule |
| | Itgax | integrin alpha X |
| | Retnlg | resistin like gamma |
| | Mmp9 | matrix metallopeptidase 9 |

| | | |
|---|---|---|
| | Ngp | neutrophilic granule protein |
| | Ltf | lactotransferrin |
| Monocytes | F13a1 | coagulation factor XIII, A1 subunit |
| | Lgals1 | lectin, galactose binding, soluble 1 |
| | Ccr2 | C-C motif chemokine receptor 2 |
| Osteoblast | Kmo | kynurenine 3-monooxygenase |
| | Bst2 | bone marrow stromal cell antigen 2 |
| | Siglech | sialic acid binding Ig-like lectin H |
| | Runx2 | runt related transcription factor 2 |
| Plasma | Jchain | immunoglobulin joining chain |
| | Rplp1 | ribosomal protein lateral stalk subunit P1 |
| | Rplp41 | Ribosomal Protein L41 |
| Pre-B Cell | Ebf1 | early B cell factor 1 |
| | Bach2 | BTB and CNC homology, basic leucine zipper transcription factor 2 |
| | Aff3 | AF4/FMR2 family, member 3 |
| | Pax5 | paired box 5 |
| | Foxp1 | forkhead box P1 |
| RBCs | Hbb-bt | hemoglobin, beta adult t chain |
| | Hbb-bs | hemoglobin, beta adult s chain |
| | Hemgn | hemogen |
| | Sox6 | SRY (sex determining region Y)-box 6 |
| T Cells | Cd3e | CD3 antigen, epsilon polypeptide |
| | Cd4 | CD4 antigen |
| | Cd8a | CD8 subunit alpha |
| | Il7r | interleukin 7 receptor |
