## Supplemental Methods for "Alzheimer’s Disease and circadian disruption sex-specifically contribute to a loss of bone maintenance in APP/PS1 model mice"

### SUPPLEMENTARY METHODS

#### Activity Monitoring

Behavior analysis used rotations per minute, aggregated in 10-minute intervals to calculate total activity, interdaily stability (IS), intradaily variability (IV), and period. Processing and calculation of behavior statistics were performed using the Actimetrics ClockLab Analysis Package (Model AM1-CA06).

#### Measurements of Aβ42 from serum and brain tissue homogenates

Tissue was submerged in 500 μL of extraction buffer (5 M guanidine hydrochloride, 50 mM Tris-HCl, pH 8.0) and homogenized using a Bead Ruptor 24 (OMNI International) with zirconium oxide beads (10 s cycle, 4 m/s, 2 cycles). Homogenates were rotated for 3 hours at 60 rpm, incubated overnight at 4oC, and centrifuged at 10,000 rpm for 20 min at 4oC. Supernatants were then diluted in 1x PBS containing 1x protease inhibitor.

#### Micro-Computed Tomography: Detailed acquisition settings

The hindlimbs were wrapped in PBS Ca-/Mg- soaked gauze and mounted in a Bruker 1272 Skyscan instrument. Male bones (LM, AP, CD, and CD/AP) were scanned at a 4 μm voxel size, with 0.5° steps through a 180° range. The instrument settings used a source voltage of 70 kV, a current of 142 μA, a 0.25 mm aluminum filter, and a 2142 ms exposure time with 2-frame averaging at a 16-bit depth. Due to scheduling limitations, female bones (LM, AP, CD, and CD/AP) were scanned in a separate batch on the same instrument after hardware maintenance. Thus, female bones were scanned at a 4 μm voxel size, with 0.5° steps through 360° range. Instrument settings used a source voltage 60 kV, a current of 166 μA, a 0.25 mm aluminum filter, and a 1166 ms exposure time with 2-frame averaging at a 16-bit depth. Scanning parameter differences required reconstruction to be conducted in two batches (males and females) to set appropriate beam hardening and segmentation thresholds. All 3D scans were reconstructed using NRecon (version 1.7.4.6, Bruker) with Gaussian smoothing kernel = 2, ring artifact correction = 10, and beam hardening correction = 15% (males) and 49% (females). Post-scan alignment correction was conducted for each sample separately.

#### Micro-Computed Tomography: Automated segmentation of trabecular volumes

Distal VOI images were thresholded between 47 and 255. In the proximal VOI, images were thresholded between 38 and 255. After thresholding, non-connected objects (speckles) in the 3D VOI were filtered and the periosteal boundary was identified by ROI-shrink wrap in 2D space (per slice), spanning holes 400 μm in diameter. The binary mask was inverted, highlighting negative space inside the periosteal perimeter. The following binary operations were used as follows to separate trabecular bridges from cortical bone and mask the internal bone volume: open (kernel: round, radius: 24 μm), dilate (kernel: round, radius 24 μm), close (kernel: round, radius: 48 μm), and erode (kernel: round, radius: 36 μm). External negative spaces due to gaps in cortical bone were removed by filtering non-connected objects in the 2D space (per slice). The resulting mask of the internal bone volume was applied to the original image, which is then thresholded and used for 2D and 3D analysis. Parameters listed here were maintained for all samples.

**Biomechanical Characterization of Bones**

In brief, the femoral head and condyles were cut using a slow-speed (150 rpm) diamond saw blade (Buehler Isomet 100, Lake Bluff, IL) to obtain a specimen with approximate cylindrical morphology. Then, a flaw, in the form of a controlled notch, was introduced at the mid-diaphyseal region on the anterior surface using the slow-speed diamond saw and sharpened using a razorblade. The mid-diaphysis of all femora was then scanned via μCT (VivaCT40, Scanco Medical AG, Basserdoft, Switzerland), using high-resolution scans acquired at 70 kVp energy, 114 mA current, 199 ms integration time, and a 10.5 mm voxel size.[1] The three-dimensional reconstruction images were used to measure the inner radius ($R_i$), outer radius ($R_o$), and notch angle with ImageJ. Samples were then tested in a three-point bending via electromechanical testing system (EnduraTEC 3200, TA Instruments, New Castle, DE) at a ramp speed of 0.001 mm/s in wet condition with a span length of 5 mm. Load and displacement curves were recorded and published equations[2–4] were used to calculate the initiation toughness (Kc, in; MPa√m), maximum toughness (Kc, max; MPa√m), and the toughening effect (ΔK; MPa√m). The stress intensity at the initiation of fracture from the notch and at maximum load during crack propagation represent the initiation and maximum toughness respectively. The toughening effect, measured as the difference between maximum and initiation toughness, was calculated to provide a measure of the energy dissipated during fracture.[4]

**Raman spectroscopy (RS) for Assessing Bone Matrix Composition**

Following fracture toughness, femora were sectioned from the mid-diaphysis using a slow speed diamond saw to produce ~500 μm transverse cross-sections. Raman analysis was performed using a Renishaw Raman Spectrometer (Ramascope 2000) equipped with a 785 nm (NIR-red) laser. A single spectrum at each morphological quadrant (N=4/sample) were acquired with 15 s of integration time per 5 accumulations. The objective was set to 50x, laser power at 25%, and a spectral range of 300-1800 $cm^{-1}$. Cosmic ray removal was performed with WiRE Project software. Baseline correction, averaging the spectrum for each individual sample, and smoothing with a 2nd order Savitzky-Golay filter was performed using a custom Matlab (R2024a) script. All peaks of interest were analyzed using peak area (PA) which represents the maximum value at a specified wavenumber. Peak area of phosphate ($\nu_1PO_4^{3-}$, ~960 $cm^{-1}$; $\nu_2PO_4^{3-}$, ~430 $cm^{-1}$), carbonate ($\nu_1CO_3^{2-}$, ~1070 $cm^{-1}$), Amide I (~1670 $cm^{-1}$), Amide III (~1242 $cm^{-1}$), methylene wag ($CH_2$-wag, 1450 $cm^{-1}$) and carboxymethyl-lysine (CML, 1150 $cm^{-1}$) were measured.

Mineral crystallinity, which serves as an indicator of crystal structure arrangement, was measured from the inverse of the full width half-maximum (FWHM) of the $\nu_1PO_4^{3-}$ peak.[5–7] The mineral-to-matrix ratio (MMR), which represents the degree of mineralization in the matrix, was calculated using both a polarization-dependent (relative ratio of $\nu_1PO_4^{3-}$ and Amide I) and polarization-independent peaks (relative ratio of $\nu_1PO_4^{3-}$ and Amide III).[5,8] The carbonate-to-phosphate ratio represents the amount of carbonate that has replaced phosphate in the crystal lattice causing a distortion in the atomic arrangement of bone. Level of Type-B carbonate substitution was measured as the ratio between the ~1070 $cm^{-1}$ carbonate subpeak and ($\nu_1$) phosphate peaks.[9,10] CML, the most abundant AGE in bone, was normalized to methylene.[11,12]

**Quantification of Fluorescent Advanced Glycation End-Products (fAGEs) in Bone Matrix**

Briefly, a ~ 2 mm section from the midshaft of each femur sample was cut (~10 mg of cortical bone). These sections were de-fatted with 100% isopropyl ether and then lyophilized overnight to remove any excess fluid. Dry samples were then placed in a glass vial, submerged with 6 N HCl, and incubated at 110°C overnight for protein extraction via acid hydrolysis. After cooling, hydrolysates were made by diluting the samples with nanopore water. These were used in two separate assays: a quinine sulphate fluorescence assay, and a collagen content assay. The first assay used a quinine stock (1 μg Quinine/mL 0.1 M $H_2SO_4$), which was diluted with sulfuric acid to create a standard curve. Triplicates of each standard and sample hydrolysate were placed in a 96-well plate, and the fluorescence was measured using a spectrophotometer (Infinite 200, Tecan Trading AG, Switzerland) at 360 nm excitation and 460 nm emission. For the second assay, the hydroxyproline content of each sample was measured. A standard curve was made from a hydroxyproline stock (2000 μg L-hydroxyproline/mL 0.001 M HCl). Solutions of chloramine-T, 3.15 M perchloric acid, and p-dimethylaminobenzaldehyde (DMAB) were added to each standard and sample hydrolysate. Each standard and sample hydrolysate were placed in a 96-well plate, and the absorbance of each sample was measured at 570 nm using the spectrophotometer.

**Bone Marrow Single Cell Analysis**

Bone marrow samples were thawed and rapidly resuspended in basal bone marrow culture media (84% DMEM-α, 15% FBS, 1% Penicillin/Streptomycin) to dilute the cryo-protectant media (44.5% DMEM-α, 44.5% FBS, 10% DMSO, 1% Penicillin/Streptomycin). The cell suspensions were prepared in 0.04% ultra-pure BSA in PBS (Ca-/Mg-) before being loaded on the 10X Genomics Chromium Next GEM controller prepared with the Next GEM 3' Single Cell v3.1 kit to generate barcoded mRNA libraries. Dual indexes (i5/i7) were added to individual mRNA libraries before PCR amplification. Subsequent clean-up was completed using SPRISelect beads. The samples were submitted to NovoGene (Sacramento, CA) for quality control, library pooling, and sequencing with paired end 150 base pair reads with 160 million reads per sample (20,000 reads per cell).

Single cell transcriptomics allows for the identification and tracking of these sensitive cell types. The Seurat FindAllMarkers function was cross-referenced with known cell type markers and the CellxGene database to annotate cells. We collected bone marrow from the femora and tibiae of male mice (8,000 cells per sample) and analyzed them using the 10X Genomics Chromium Next platform. We sequenced at a depth to ensure at least 20,000 reads per cell. Following sequencing (Novogene), the raw data was passed to the 10X Genomics cloud analysis platform, before downstream analysis in the Seurat package (version 5). The Scanpro package was used to simulate random samples of cell populations. We used 500 iterations, verified by stabilization of Z-score statistics. The initial exploration of the bone marrow transcriptome compared bulk cell expression with the Likelihood Ratio Test (LRT) implemented in the Seurat FindMarkers function. Cells were clustered with the Seurat implementation of K nearest neighbors.

**Proteomics**

Batch effect and imputation corrections were made using the LIMBR pipeline[13], accounting for genotype and lighting conditions separately. Proteins with more than 30% missing values were excluded, remaining missing values were imputed using a K-nearest neighbors' algorithm (K = 10). Expression level was quantified as the average peptide signal-to-noise ratio per protein.
ECHO fitting was applied separately to each condition using the free-run paired mode with smoothing, normalization, and linear detrending.

**Supplementary Methods References**
